# Reframing enzyme function prediction as conditional generation

**DOI:** 10.64898/2026.09.09.750355

**Authors:** William JF Rieger, Sebastian Häussermann, Luca Herrmann, Zecheng Li, Béla P. Frohn, Manuel Maluenda, Gabriela Lobinska, Sahil Loomba, Julia C. Reisenbauer, Mikael Bodén, Alexander Tong, Ariane Mora

## Abstract

Enzymes frequently exhibit promiscuous activity beyond their native roles, providing starting-points for new functions. Finding these promiscuous enzymes, especially for non-native chemical transformations, is challenging but highly valuable, as they promise novel, sustainable solutions for chemistry and biotechnology. However, current machine learning methods are poorly suited to discovering unseen chemistry as they often frame function prediction as closed set classification or a retrieval task. Here, we present *Fluxion*, a generative deep learning framework that learns enzymatic catalysis by modeling dynamic electron flow trajectories across the enzyme’s catalytic residues. By combining both synthetic chemistry and biochemical datasets with protein language model representations, Fluxion generates multi-step electron-flow trajectories analogous to the arrow-pushing representations used to describe enzyme reaction mechanisms. Generation is conditioned on enzyme context, including the enzyme sequence, catalytic residues, substrates, and cofactors. We show that this conditioning allows Fluxion to learn enzyme-dependent regioselectivity across cytochrome P450 enzymes with different sequences shifting the predicted reaction sites for the same substrate. We then demonstrate that Fluxion’s embeddings are useful for downstream tasks, such as specificity prediction on two experimental datasets, with and without finetuning. Finally, we show that Fluxion has the potential to transfer synthetic chemical logic to biology; it can generate the observed non-native product from real-world non-native directed evolution screens. Our results establish a proof of concept that generative modeling through mechanistic representations of enzymes can shift enzyme function prediction beyond static database retrieval and closed set classification to function generation. This conceptual framework provides a stepping stone towards an in silico generative method to discover non-native biocatalysts.

## 1 Introduction

Directed evolution allows us to repurpose and optimize enzymes to effectively catalyze promiscuous non-native reactions, by taking enzymatic starting points that exhibit low levels of activity toward a desired function and engineering them toward targeted biocatalytic tasks [1, 2]. Yet discovery of these starting points remains a bottleneck in biocatalysis [3].

Most state-of-the-art machine learning (ML) approaches treat enzyme function as a retrieval problem, typically assigning enzymes to a closed set of Enzyme Commission (EC) numbers, an annotation that does not necessarily indicate specific reaction or extend to non-native reactions [4–6]. Recently, retrieval for specific reactions has also been attempted [7], which are either limited to a set of reactions in the database or aim to predict a binary activity between substrates and enzymes. Methods developed to predict specificity such as ESP [8] and EzSpecificity [4], are unable to generalize to novel chemistry [9]. Furthermore, these methods rely on binary classification of enzyme-substrate pairs rather than generating complete reaction or product distributions. As retrieval based methods are constrained by the database used expanding biocatalysis into the non-native chemical space remains out of reach. Consequently, state-of-the-art ML enzyme-reaction prediction models fail to generalize to reactions not observed during training [9, 10].

Generative methods use ML to learn the probability distribution of complex data, which can then be conditioned to sample from a specific subspace. By approximating the distribution, generative models can create plausible, previously unseen configurations rather than being restricted to reproducing or retrieving examples from the training set. They have shown success in generalization across domains in life-sciences, including de novo protein design [11], enzyme design [12, 13] and small molecule design [14]. Most directly relevant to the present work, Joung et al. [15] introduced FlowER, extending generative modelling to chemical reactivity by representing reaction mechanisms as trajectories of electron redistribution. Despite progress in other domains, predicting enzyme function based on generative modeling is nascent. For example, auto-regressive models aim to learn the syntax of enzymes by trying to predict the product as the next part of a sentence (e.g., letter in a SMILES string), however, this approach is limited often conditioning enzyme identity through categorical labels rather than molecular representations of the enzyme itself [16]. More recently, EzSolver finetuned FlowER with catalytic residues to predict polar enzymatic mechanisms [17]. Consequently, these models can associate enzyme classes with reaction outcomes, but cannot directly learn how variation in enzyme sequence or structure gives rise to differences in biochemical function.

Enzymatic catalysis is governed by the enzyme’s macromolecular environment where active-site catalytic residues are necessary but insufficient alone to determine reactivity. To capture non-native and promiscuous function beyond the natural domain, a generative model needs to overcome three challenges: first, it must learn how catalytic residues coordinate multi-step reactions; second, it must incorporate global context; and third, it must overcome the limited scale of biochemical reaction data by learning transferable chemical reactivity from much larger synthetic chemistry datasets, effectively bridging the domains of synthetic organic chemistry and biochemistry. To address these requirements, we developed Fluxion, a generative deep learning framework that models biocatalytic reactions as stochastic electron-flow trajectories between reactant and product states using a Schrödinger bridge formulation adapted from Liu et al. [18]. Fluxion is pretrained on synthetic organic chemistry data and subsequently fine-tuned on biochemical reaction data. To enable the model to learn from the whole enzyme context rather than just catalytic residues, we condition generation on protein language model (PLM) embeddings. The stochastic nature of Fluxion’s bridge model enables the sampling of potential promiscuous enzyme activity for both natural and non-native reactions. We demonstrate that generative models like Fluxion enable zero-shot prediction of complex reaction mechanisms and can capture non-native biocatalytic functions.

## 2 Results and Discussion

### 2.1 Modeling of enzymatic reactions to generate products

We formulate enzyme function prediction as a generative task. Conceptually, Fluxion must learn two complementary aspects of enzymatic reactivity: which electron-flow trajectories are chemically plausible which we do similarly to Joung et al. [15], but also, how the surrounding enzyme context changes the relative likelihood of those trajectories. Fluxion generates single- and multi-step reaction flows conditioned on both catalytic residue representations and global protein sequence context (Fig. 1A). The model uses substrate and cofactor SMILES strings, and user defined “residues of interest” (catalytic and binding site annotations) (Fig. 1B), all represented by the bond-electron (BE) matrix formalism [21] as in FlowER for stepwise mechanism generation [15]. As catalytic residues are insufficient to define the selectivity and specificity of an enzymatic reaction, we use cross attention to integrate the overall enzyme context. This is achieved by including the sequence and per-token embeddings from the protein language models ESM2-3B as an input [22], enabling Fluxion to learn from both the chemical mechanisms (USPTO, M-CSA) and enzyme context (EnzymeMap) (Fig. 1C). During inference, Fluxion’s bridge model draws *N* generative samples, producing a frequency-ranked product distribution filtered for chemical validity and cofactor preservation. Rather than returning a single reaction assignment, this sampling procedure represents alternative chemically plausible outcomes within a conditional distribution, allowing candidate products or reaction steps to be ranked by their frequency among generated trajectories.

**Figure 1:**
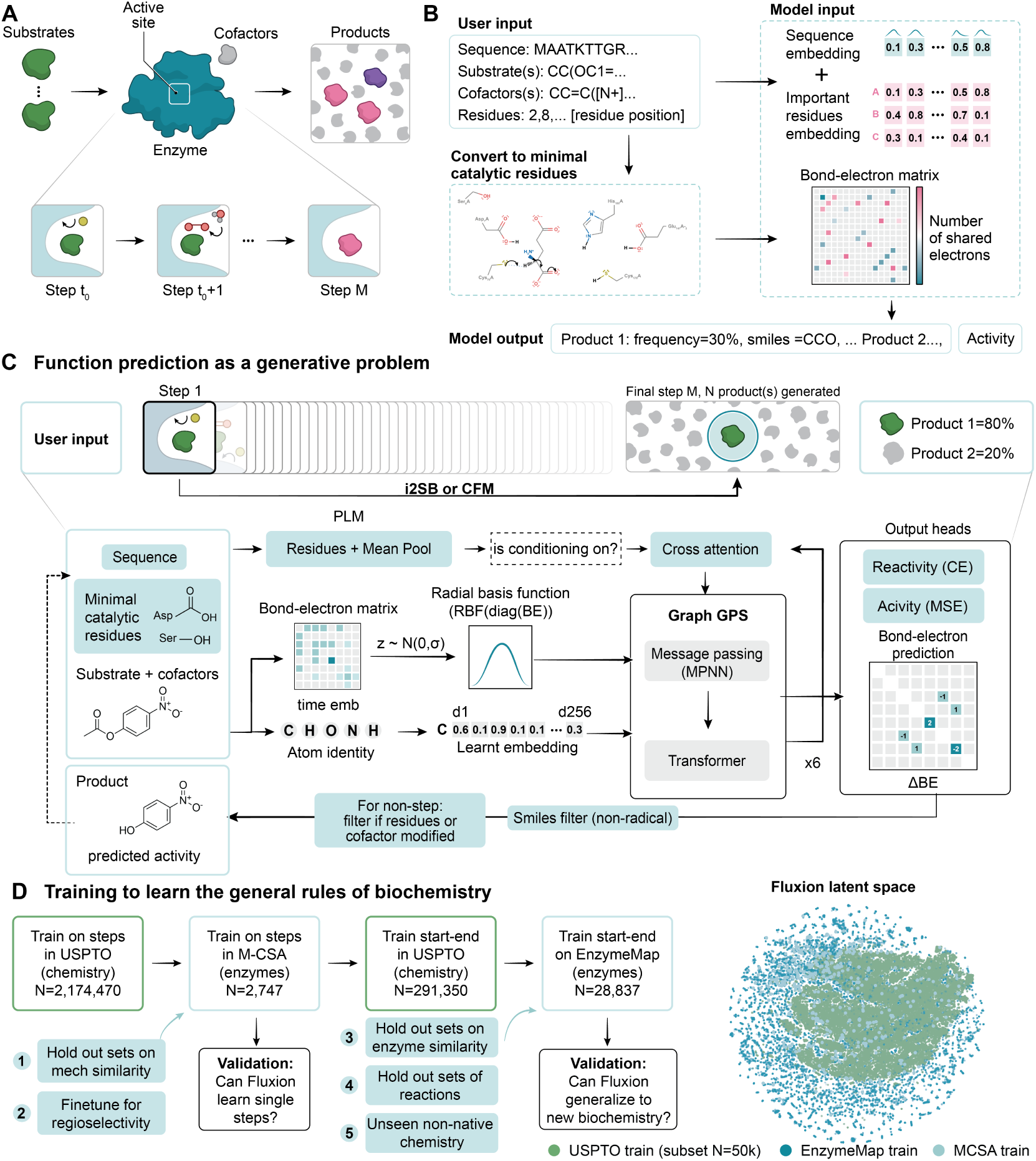
Fluxion: a model to generate enzyme products and stepwise mechanisms. **A**) Fluxion takes substrates, cofactors, and enzyme sequence information and generates possible products and reactivity. **B**) User workflow: inputs (SMILES, sequence, and residue positions) are automatically processed, returning frequency-ranked product distributions across a user specified number of generated samples along with reactivity. **C**) Fluxion predicts single- or multi-step trajectories by integrating Bond-Electron (BE) matrices (substrates, cofactors, catalytic residues) with ESM2 protein embeddings via cross-attention, employing optional SMILES chemical correctness filters for non-radical reactions. Fluxion primarily outputs a set of generated product SMILES. In analyses of enzyme reactivity and activity, we additionally use optional auxiliary heads that predict whether an enzyme-substrate pair is reactive (binary 0/1) and, for reactive pairs, a continuous activity score between 0 and 1. These auxiliary outputs are separate from product generation and provide additional signals for ranking enzyme-substrate outcomes. **D**) Evaluation framework for cross-domain synthetic and biochemical performance on held-out experimental datasets. T-SNE embeddings from Fluxion’s encoder.

We evaluate the capabilities of this framework through four experiments (Fig. 1D). (1) We first ask whether Fluxion learns transferable chemical reactivity by evaluating recovery of held-out enzymatic mechanisms across increasingly dissimilar reactions (Fig. 2). (2) We then test whether protein context can shift the learned distribution towards enzyme-specific outcomes, using cytochrome P450 regioselectivity as a benchmark (Fig. 3). (3) We next evaluate whether the same framework can support prediction of enzyme-substrate reactivity and activity across experimental screening datasets (Fig. 4). (4) Finally, we ask whether chemical knowledge learned across synthetic and bio-chemical reaction space can extend beyond native enzyme function by testing product generation for experimentally validated non-native directed-evolution campaigns (Fig. 5).

**Figure 2:**
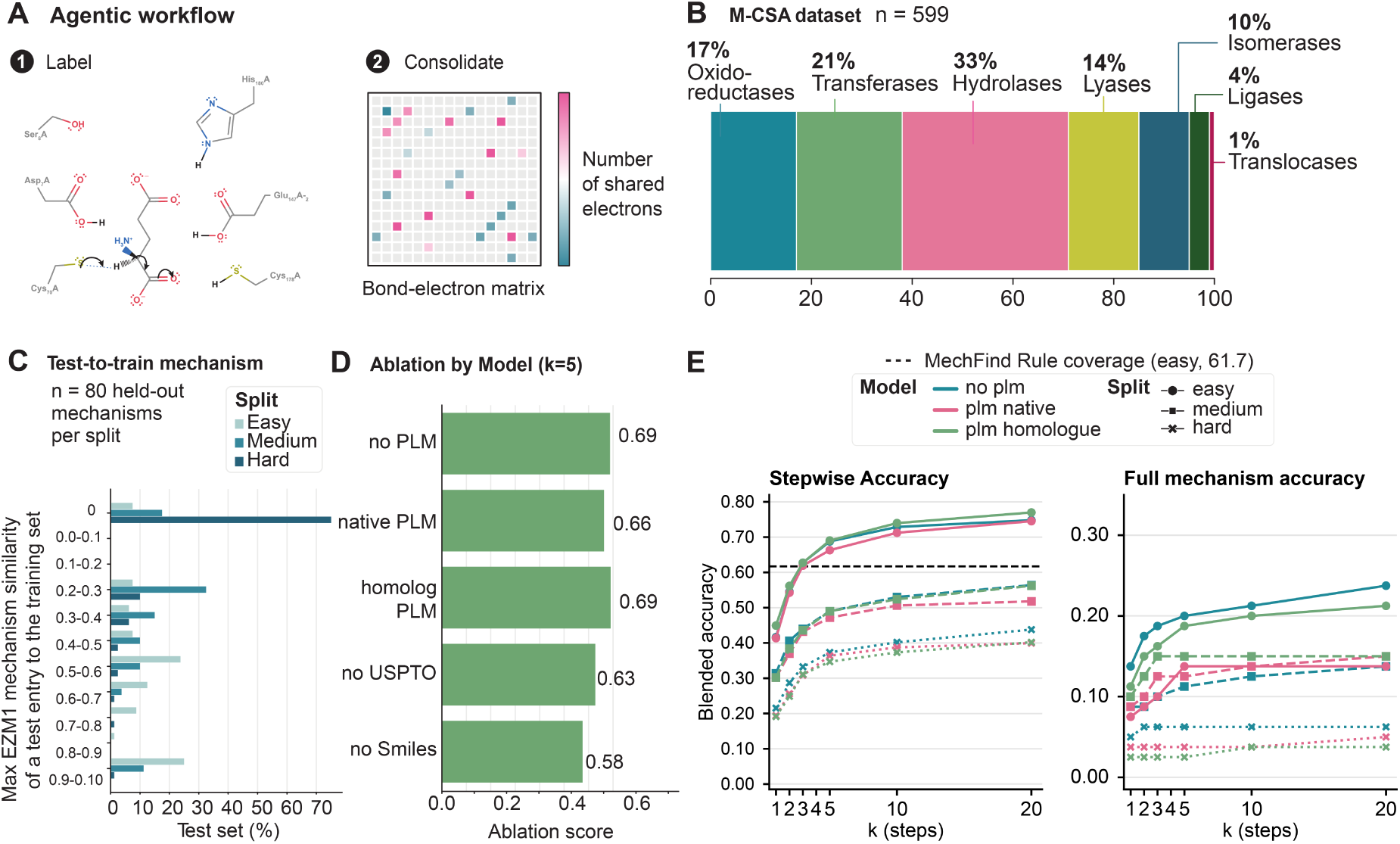
Performance of Fluxion for enzyme mechanism prediction. **A**) Illustrative mechanistic step from M-CSA entry 1. M-CSA mechanistic annotations were processed using an agentic workflow to atom-map reactions and standardize inconsistencies in representation across entries. **B**) Distribution of the 599 reconstructed reactions across EC classes, showing substantial class imbalance, with hydrolases (EC 3) comprising 33% of the dataset. **C**) Easy, medium and hard benchmark splits were constructed using the Jaccard-based pairwise mechanism similarity of Ribeiro et al. [19], with progressively lower similarity to mechanisms in the training data. The hard set predominantly contains reactions with zero observed similarity and was further supplemented with ten reactions selected for long mechanisms or large total number of atoms (Fig. S2). **D**) Top-5 stepwise accuracy across Fluxion ablations, assessing the contribution of PLM conditioning, USPTO pretraining and SMILES conditioning. **E**) Top-k stepwise and pathway accuracy across the easy, medium, and hard benchmark splits. Stepwise accuracy measures recovery of the correct mechanistic step within the top-k predictions, while pathway accuracy requires recovery of every step in the complete mechanism within the top-k predictions per step. The dashed line indicates rule coverage for MechFind [20] when regenerating its rule set using the easy training data.

**Figure 3:**
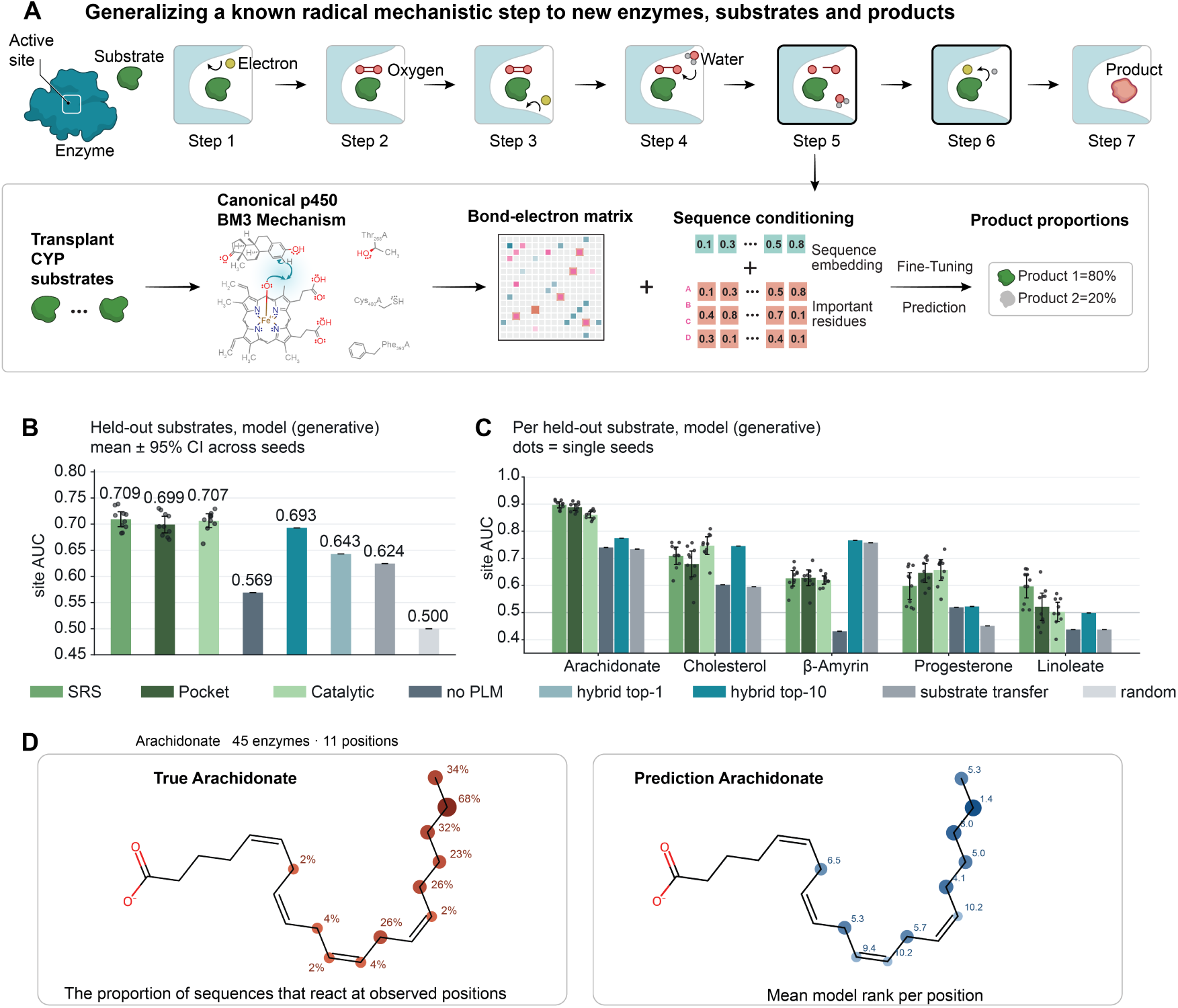
Fluxion can learn a regioselective step in P450-BM3 biocatalysis. **A**) Mechanistic steps of the CYP aliphatic hydroxylation of P450-BM3 enzymes (M-CSA entry 699) [26]. Fluxion is fine-tuned using publicly available CYP reaction data to generate sequence-conditioned distributions over alternative selectivity-determining hydrogen atom transfers. **B**) The resulting product probabilities are used to rank candidate hydroxylation sites across five held-out substrates. Three sequence conditioning variants of Fluxion are compared with hybrid sequence/substrate-transfer baselines, a substrate-transfer baseline, and random ranking. SRS represents the model conditioned with additional ESM2 tokens for residues identified to be important for substrate recognition. Pocket conditioning further subsets SRS sites to those within the binding pocket. The catalytic model is conditioned only on catalytic residue tokens. The noPLM model uses no PLM conditioning. Site AUC is the mean probability across enzyme-substrate pairs that an observed site outranks an unobserved site (ties = 0.5). **C**) Site AUC for each of the five held-out substrates. **D**) Experimentally observed and predicted arachidonate hydroxylation patterns across 45 CYP enzymes. The observed distribution (left, red) shows the percentage of enzymes that hydroxylate each substrate position. The predicted distribution (right, blue) shows the mean rank assigned to each position across the corresponding enzyme-specific predictions, where a lower rank indicates a more highly predicted site (site rankings and observed positions for all sequence-substrate pairs in Figs. S5–S9).

**Figure 4:**
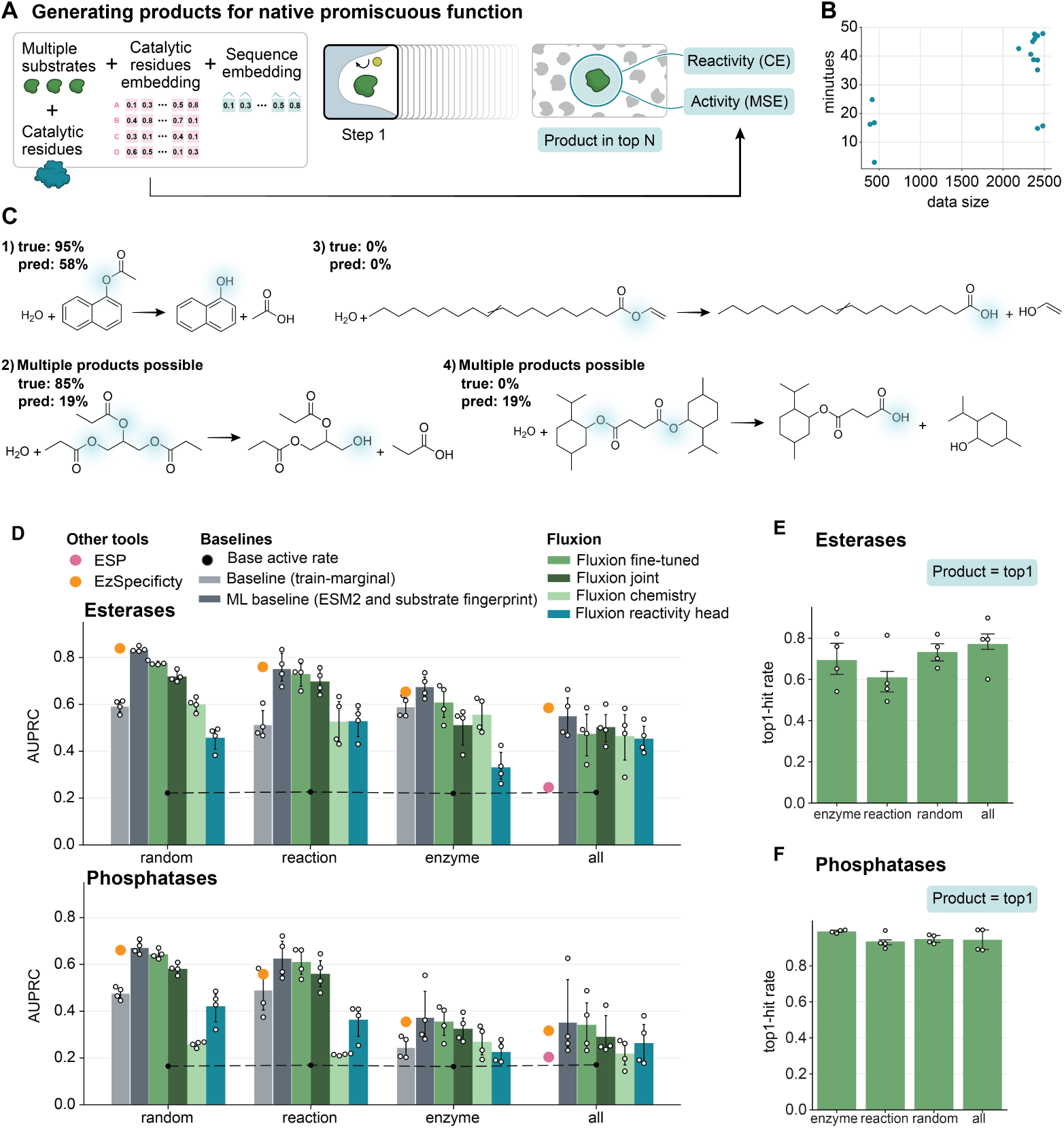
Evaluation of performance on native substrate selectivity and native product generation. **A**) To learn native products rather than single steps, Fluxion was trained to generate a final product, given an input sequence, the catalytic residues, substrates and cofactors. Products are generated in a single step. **B**) The time taken to “virtually” screen over 2,000 enzyme-substrate pairs is less than 1 hour, generating 8 samples per reaction-enzyme pair. **C**) Using the sets as defined in EzSpecificity for training and testing [4], we evaluate the performance of the reactivity head as the reaction probability using Area Under the Precision-Recall Curve (AUPRC). The training sets were used to finetune Fluxion for each dataset, learning a “reactivity” head, results for reactivity prediction vs the baseline activity of a substrate are shown here. **D**) Evaluation of splits from [4]. The Base active rate is the rate of positive samples, while the baseline is the marginal for the observed training data, for the enzyme it is the marginal of the substrate (e.g., the reactivity of the substrate), and for the reaction it is the marginal of the enzyme (e.g., how promiscuous is the enzyme). As we use the marginal of the observation in the training set when both are held out (in “all”) there is no baseline. Data for EzSpecificity and ESP are as reported in EzSpecificity’s main Figure 5F (all) and EzSpecificity’s Table S5 (enzyme, reaction, random) using the same splits [4]. Additionally we train a Gradient Boosting Model (GBM) on ESM2 embeddings and Morgan fingerprints as a ML baseline, to compare the embeddings from Fluxion, where embeddings are taken from: Fluxion is pretrained on chemistry (Fluxion chemistry) and biochemical reaction data (Fluxion joint) then fine-tuned on publicly available enzyme activity data for esterases and phosphatases (Fluxion finetuned) which also includes a reactivity head (Fluxion reactivity head). **E**) Despite training two new heads, to learn reactivity and activity, the model retains high product generation capacity across the splits, even for unseen substrates and enzymes in esterases, and **F**) phosphatases.

**Figure 5:**
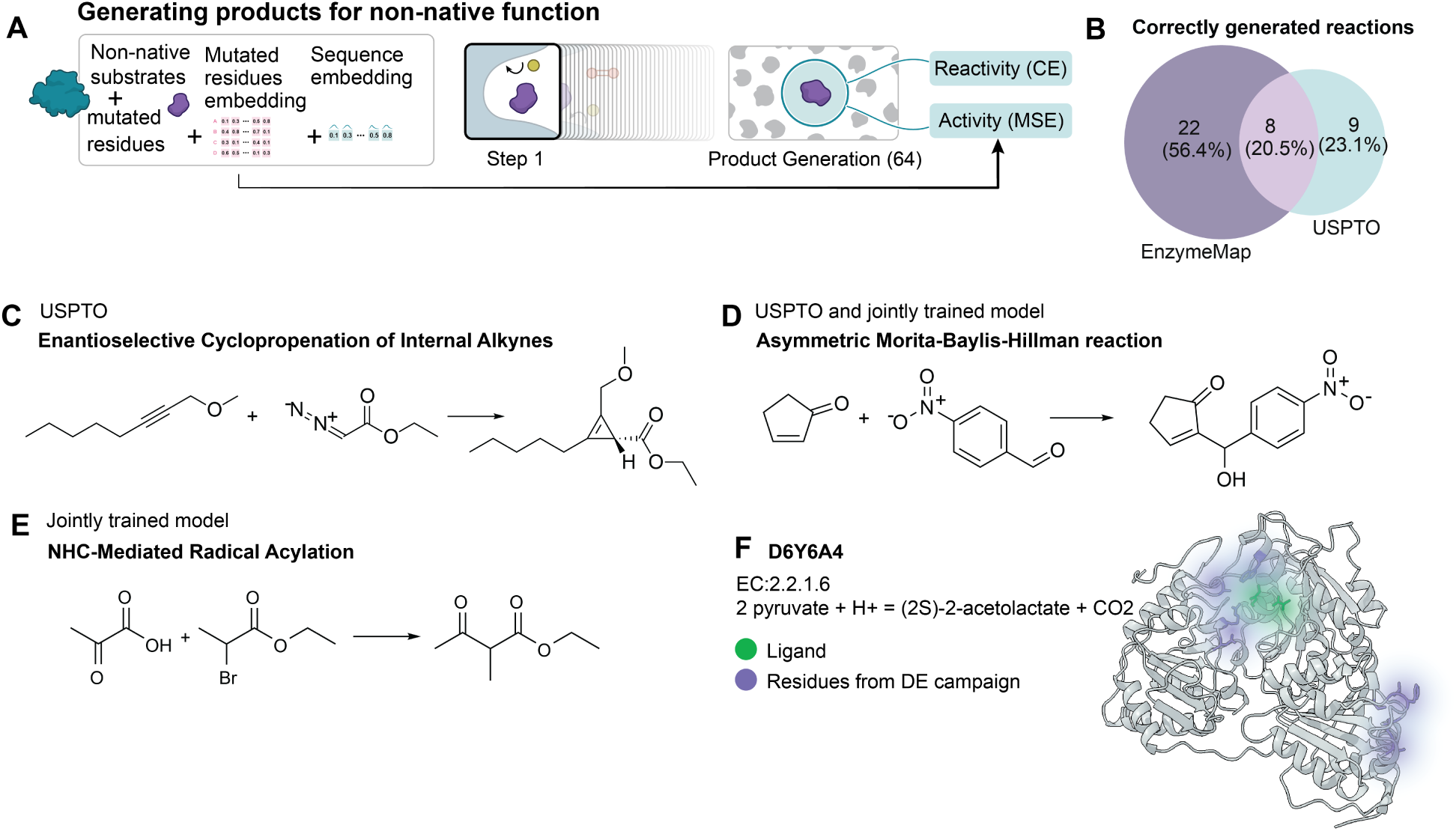
Transfer from the synthetic chemical to biochemical space. **A**) Here we evaluate the performance of the single step start-end (substrates to products) Fluxion model, and evaluate the performance of the model trained using either USPTO (chemistry only) or USPTO + EnzymeMap (both chemistry and biochemistry). **B**) For the tested reactions that had a correct product generated, we show which reactions were predicted by only one model, or both, showing that some reactions are jointly predicted, while others require the training data from both models. **C**) An example of a reaction detected only with the Fluxion model trained on USPTO (only trained on chemistry), from Chen and Arnold [35]. **D**) Both versions of Fluxion (trained independently on chemistry (USPTO) or with enzyme data (joint e.g. USPTO + EnzymeMap), were able to detect this reaction), from Wang et al. [36] **E**) Only the jointly trained model was able to predict that Thiamine- and Flavin Dependent Enzymes could perform NHC-Mediated Radical Acylation [37]. **F**) taking the table from the initial screening campaign in the SI of the paper from Kato et al. [37], we tested whether a priori the model could have been used to facilitate manual selection, finding that the majority of the enzymes were correctly predicted to have activity, here we folded the best enzyme from the campaign with the co-folding model Chai [38], taking the highest scoring result (0.3025) of 5 samples, showing the native function of this enzyme.

### 2.2 Performance on the task of enzymatic mechanism generation

Bond-Electron (BE) matrices [21] provide a data format for ML models to predict single-step organic reaction mechanisms [15, 23–25], but converting biological databases like M-CSA [26] to BE format is non-trivial, challenged by atom-labeling inconsistencies across independently drawn steps. To construct a dataset suitable for our model, we developed an agentic annotation and reconstruction workflow that converts the stepwise mechanisms in M-CSA into bond-electron (BE) matrices (Fig. 2A). Of the 734 M-CSA mechanistic entries, 599 (81.6%) were successfully reconstructed, with 14% containing one or more warning-flagged steps requiring additional caution during interpretation (S. Tables 1-2; Methods). The resulting dataset spans the major enzyme classes represented in the M-CSA (Fig. 2B), including oxidoreductases, transferases, hydrolases, lyases, isomerases, ligases, and translocases, and encompasses mechanistically diverse transformations including both polar and radical chemistry. We next constructed three test-to-training splits designed to evaluate increasingly out-of-distribution mechanism predictions (Fig. 2C). Test mechanisms were assigned according to their maximum mechanistic similarity, a metric developed by Ribeiro et al. [19], to any training example, producing easy, medium, and hard splits with progressively lower similarity to the training set (see Methods). Each split contained 80 held-out mechanisms, with varying numbers of total steps. The hard split had a mean mechanism similarity of 10.3% to the closest training mechanism, whereas the easy split retained close mechanistic analogues with a mean of 61.4% similarity.

Fluxion maintained single-step predictive performance across increasingly difficult mechanism-similarity splits, including predictions beyond the coverage of training-derived mechanistic rules (Fig. 2E; performance by EC class in Fig. S4), namely, a top-1 stepwise accuracy of 45%, decreasing to 32% and 22% on the medium and hard splits, respectively. Accuracy increased with k across all three splits, indicating that the correct elementary transformation was frequently retained among the model’s predictions even when it was not ranked first. For comparison, the maximum stepwise rule coverage of training-derived MechFind [20] rules were 61.7%, 45.0%, and 32.0% for the easy, medium, and hard splits, respectively, representing the upper bound that MechFind can predict. Fluxion exceeded these values at k=3 across all three benchmark difficulties, indicating that the model can recover transformations that were not directly represented by the corresponding rule set.

We next assessed the performance of Fluxion on complete reaction mechanism predictions rather than individual stepwise performance. Reaction mechanisms for the held-out test set were generated iteratively by using beam search and ranked according to their length-normalised step likelihoods. Pathway-level accuracy was substantially lower than stepwise accuracy and declined with increasing benchmark difficulty: at k=5, product-informed pathway accuracy was 20.0%, 11.3%, and 6.3% on the easy, medium, and hard splits, respectively (Fig. 2E). The maximum full-mechanism coverage (upper bound on exact mechanism accuracy) of training-derived MechFind rules similarly decreased from 46.0% on the easy split to 22.5% and 6.3% on the medium and hard splits, highlighting the limited representation of complete mechanisms in the more difficult test sets. Thus, while Fluxion retained the capacity to identify individual transformations for mechanisms without close training analogues, reconstruction of complete pathways remained increasingly challenging with mechanistic distance from the training data.

We attempted to fine-tune FlowER on M-CSA as a point of comparison with an alternative generative architecture built on the same BE-matrix formulation. However, fine-tuning did not converge, collapsing to the no-displacement terminal state at every step, and its preprocessing pipeline rejected 20.7% of easy-split steps, so the two models could not be evaluated over an identical set of steps (Methods). We therefore report no fine-tuned FlowER baseline. Concurrent work [17] successfully fine-tunes FlowER on a polar subset of M-CSA reactions and introduces EZSolver, which combines evaluator-guided bidirectional beam search with chemical and 2D structural constraints to improve the consistency of generated mechanisms. Adding comparable trajectory scoring and filtering to Fluxion is a promising route to improving complete mechanistic pathway prediction.

PLM conditioning provided little additional benefit for mechanism recovery (Fig. 2D), consistent with the limited sequence redundancy in M-CSA, where most entries lack a detectable homologue within the dataset (Fig. S3). The M-CSA therefore provides little supervision for learning how sequence variation shifts mechanistic outcomes. In contrast, USPTO pretraining improved performance, supporting the value of broad chemical pretraining for this task. We retain PLM conditioning to enable subsequent fine-tuning in settings where protein context is discriminative, as exemplified by the following enzyme-specific tasks.

### 2.3 Learning enzyme-dependent regioselectivity with a generative reaction model

The ability to control chemo- and regioselectivity is essential to chemical synthesis, and enzyme engineering has the potential to provide scalable solutions for this task. Cytochrome P450 enzymes (CYPs) are a particularly useful system for studying regioselectivity because closely related enzymes can catalyse the same reaction at different substrate positions, as e.g. the hydroxylation of various C H bonds. Computational models of CYP reactivity generally formulate regioselectivity as atom- or bond-level classification/ranking [27–30]. A generative reaction model offers an alternative approach in which alternative regioselective outcomes are represented within a conditional distribution over possible reaction steps (Fig. 3A). We therefore asked whether Fluxion could learn enzyme-dependent distributions of CYP regioselectivity, providing a proof of concept for this formulation. We hypothesized that conditioning the generative model on enzyme sequence would shift probability mass towards the regioselective outcomes observed for individual CYPs. To test this, we derived a new benchmark from the CYP substrate-promiscuity dataset of Mahood et al. [9] and fine-tuned Fluxion’s mechanistic step model to generate regioselective outcomes.

The benchmark from Mahood et al. [9] evaluates whether an enzyme accepts a given substrate while for our task we test whether the model can identify the specific site of a substrate that is activated by a specific CYP enzyme. All aliphatic hydroxylation reactions involving five substrates with high regioselective diversity, arachidonate, cholesterol, *β*-amyrin, progesterone and linoleate, were held out from the training set (see Methods). Each substrate was associated with 20 to 45 active CYPs and hydroxylation was observed across 5 to 11 distinct C H positions, providing substantial variation between enzymes. Enzymes reacting with arachidonate showed particularly broad regioselective promiscuity, with individual CYPs frequently hydroxylating multiple positions (Fig. 3D).

We effectively demonstrate that Fluxion’s generated regioselective outcomes are dependent on enzyme sequence, rather than merely reproducing substrate-specific reaction priors. Across the five held-out substrates, Fluxion achieved a mean overall site-ranking AUC of 0.709 (0.68-0.74 across ten training replicates), exceeding models without protein sequence conditioning and substrate-transfer baselines that predict reactive sites by transferring observations from chemically similar training substrates (Fig. 3B). On arachidonate, Fluxion achieved a mean AUC of 0.897, outperforming a test-set-informed substrate prior (0.811) constructed from the true hydroxylation sites of other test enzymes acting on the same substrate (Fig. 3C; site rankings and observed positions in Figs. S5–S9). These results indicate that Fluxion captures enzyme-specific information beyond the population-level regioselective preferences of enzymes that react with arachidonate.

Much of the sequence dependent signal could nevertheless be recovered from enzyme homology. A hybrid baseline combining substrate specific reactivity with sequence similarity performed marginally below Fluxion, with an overall site-ranking AUC of 0.693 (top-10 hybrid), indicating that a substantial fraction of the predictive information encoded by the ESM2 representation correlates with evolutionary relatedness (Fig. 3B). Fluxion retained a modest improvement over this baseline, with the exception of *β*-amyrin, suggesting that the learned representation captures additional information relevant to regioselectivity beyond global sequence similarity. To test whether the selection of additional residues as compared with only the catalytic residues improved regioselectivity prediction, we independently fine-tuned three Fluxion variants conditioned on ESM2 embeddings from different CYP residue sets: canonical substrate-recognition sites (SRS), a reduced binding-pocket subset, or catalytic residues only (Methods; Fig. 3B). Performance was similar across all three splits indicating that the per-token-embeddings were not adding additional relevant information.

Together, these results provide a proof of concept that enzyme-dependent regioselectivity can be represented within the outcome distribution of a generative reaction model. However, the strong performance of the homology baseline and limited benefit of alternative sequence conditioning suggests that learning regioselectivity from sequence alone remains challenging. Future models incorporating structural information about enzyme-substrate interactions may improve these results.

### 2.4 Fluxion learns native enzyme promiscuity

Directed evolution (DE) has enabled the effective optimization of biocatalysts toward non-native transformations [1, 2]; however, identifying initial promiscuous starting variants that can later be optimized still largely relies on experimental screening and chemical intuition. To assess whether generative architectures or embeddings from generative models can be applied to this problem, including their current capabilities and failure modes, we evaluated Fluxion on both native, and non-native enzyme product generation, reactivity and activity prediction.

Analogous to confidence heads in structural models such as AlphaFold [31], we posited that Fluxion could jointly learn two auxiliary targets: a reactivity head, which predicts whether a given enzyme and substrate will react, and an activity head, which estimates the expected level of catalytic activity when a reaction occurs. Additionally, the generative component was retrained to predict the product directly from the reactants in a single step, rather than predicting individual mechanistic steps (Fig. 4A), making it efficient to screen across many enzyme-reaction pairs (Fig. 4B). We evaluated this architecture on experimental data from two enzyme screening campaigns in which activity was profiled across esterases [32] and phosphatases [33] preprocessed by Goldman et al. [34]. The esterase benchmark spans 146 enzymes tested against 93 substrates from Martínez-Martínez et al. [32] and the phosphatase dataset is 218 enzymes by 92 substrates [33], once filtered, see Methods. The datasets display varied reactivity profiles ranging from universally accepted substrates to those showing no conversion across any tested enzymes (Fig. 4C). A single enzyme can also produce multiple distinct product outcomes.

Using the benchmark splits established by Cui et al. [4], we found that a simple gradient boosting model (GBM) on Fluxion’s embeddings (Fig. 4D), performs on par with the SOTA model EzSpecificity which leverages docked structures to learn specificity between enzymes and substrates (Cui et al. [4]), and better than sequence based model ESP [8], however, since some reactions were dropped owing to atom balancing, this can be considered indicative rather than concrete. The same GBM model, trained only on substrate (Morgan fingerprint) and enzyme embeddings (ESM2) performs on-par with both Fluxion’s embeddings and EzSpecificity’s model suggesting that a simple model using reasonable representations performs similarly to more complex models (Fig. 4D). The learnt reactivity head in Fluxion was unable to learn prediction at a comparable rate with the GBM, however, the fine-tuned model with the reactivity head generally outperforms the base Fluxion model (Fluxion joint) (Fig. 4D). In summary, the learnt embeddings for chemistry appear as informative for this downstream task as those of baseline approaches, yet do not appear to add any new information, see SI (S18).

We next tested whether Fluxion can accurately prioritize candidate enzymes for a specified, unseen substrate, i.e. predict highly active vs lowly active enzymes using the activity head. The average correlation is weakly positive for esterases at *ρ* = 0.32 p *<* 0.05 (Fig. S10), while no global activity signal could be learnt across the phosphatase dataset, (Fig. S12), or from the negative control e.g. random sampling (Fig. S13). Across both datasets the accuracy of the activity head varies with substrates (Figs. S14-S17). Finally, the addition of auxiliary prediction heads did not compromise the underlying generative framework, and product generation retained high fidelity across all benchmark splits, with only 8 generative samples (Fig. 4E-F). These results suggest that the generative framework can be used for specificity prediction, however, would likely benefit from the inclusion of other features, such as structure. Comparable performance was maintained when pre-training exclusively on synthetic reactions from the USPTO corpus (Fig. S11), demonstrating that large-scale pre-training on general chemical reactions could provide a viable route toward generalizable biocatalytic modeling.

### 2.5 Generating non-native function

Since Fluxion has been trained on both organic synthetic chemistry and biochemistry, we hypothesized it could be used beyond native function prediction to identify non-native function. Using 206 experimental enzyme engineering campaigns from Long et al. [39], we evaluated whether Fluxion can generate non-native products for a specific enzyme, substrate, and cofactor combinations (Fig. 5A). Enzymes passed to the model included both the initial evolutionary variants that unlocked detectable promiscuous activity and the final engineered variants obtained after extensive DE optimization campaigns. Using a budget of N=64 generated samples, the jointly trained Fluxion model generated the correct non-native product for 16% of evaluated reactions (N=33/206) (Fig. 5 B-E). We looked at the initial screening campaign from one experiment, (exp LLM83), NHC-Mediated Radical Acylation Catalyzed by Thiamine- and Flavin-Dependent Enzymes, from Kato et al. [37], and ran their enzyme screen for initial activity through Fluxion. In this campaign predictions varied across the 14 screened enzymes (11 active, 3 inactive calls), suggesting the model is not producing a substrate-determined constant (Fig. 5E-F). To evaluate whether this holds across the EnzEngDB, we tested whether randomly swapping the enzyme impacted non-native product generation. When running the 206 reactions, we find that generation capacity is not dependent on the enzyme: for randomly selected enzymes from the EnzEngDB we see no difference in the number of correctly generated products within 64 samples (N=35/206), in fact a slight increase when randomly perturbing the sequence. As EnzEngDB is dominated by heme-dependent enzymes, we hypothesize that this result highlights the contribution of the cofactor towards the chemistry. These results for the baseline Fluxion model are unsurprising given the absence of negative examples in training. Fluxion never observes enzyme-reaction pairs labelled inactive, and so receives no signal for when not to generate a product. Hence, applying Fluxion to enzyme screening will require fine-tuning on experimental negatives, or hard negative imputation based on shared cofactors and function.

When considering the model trained only on chemistry, different products were successfully generated, indicating that the reactions that are further from native function may be weakened by the post-training on the biochemistry dataset, (Fig. 5C, Fig. S19). We then sought to test whether the model was just recapitulating the training data, or had learnt unseen product generation. When testing the Tanimoto similarity to the training sets we found that both the hits and misses followed a similar distribution (*µ*=0.79 for hits, vs *µ*=0.72 for misses for EnzymeMap, and *µ*=0.78 for hits and *µ*=0.79 for misses for USPTO). While several of the TrpB reactions were in the training set, the other 22 identified reactions are genuine hold outs suggesting that Fluxion has the capacity to learn to generate the correct unseen products for non-native reactions.

## 3 Conclusion

Biocatalysis provides a sustainable alternative to traditional chemical synthesis, yet exploiting enzyme promiscuity for non-native reactions still largely relies on experimental screening to identify initial active variants. With Fluxion we explored the use of generative models for the task of enzyme function prediction. We found that generative models can effectively learn single steps for enzymes as in concurrent work [17], and that in this task, enzyme conditioning presents no benefit, likely because of the lack of sequence redundancy in the training data. However, by designing the model architecture to take in conditioning, we can apply the model to downstream enzyme function prediction tasks where they are useful.

First, we finetuned the single step Fluxion model to learn regioselective generation for a challenging radical reaction across cytochrome P450s. On this task, Fluxion performs comparatively to homology and substrate transfer while maintaining generation fidelity, suggesting that with sufficient data, generative models can learn the nuances of enzyme catalysis. We then asked what Fluxion’s reactant to product model encodes, finding that the learned embeddings are useful for the downstream task of specificity prediction, meeting the ML baseline using reaction fingerprints and ESM2. This suggests that the model learns a useful biochemical representation, but that more information would need to be included (e.g. all individual steps, or structure) to overcome the baseline performance. Lastly, we posited generative models could be used for non-native discovery, and found that while Fluxion can generate non-native function, the model is unable to distinguish between enzymes across this task. Hence, without fine-tuning, generation is driven by substrate and cofactor chemistry rather than enzyme identity. We attribute this to the curation of the training data, which contains little sequence redundancy and no negative examples, leaving no contrastive signal between related enzymes, and none for when not to generate a product. Thus, several challenges remain before this framework can be applied prospectively. A defining feature of Fluxion is its atom-conserving formulation, as no new atoms can be created or removed during the generation process, users must specify all reaction participants that contribute atoms to the products, including relevant cofactors or co-substrates in the input. While this constraint imposes additional requirements on reaction representation and data curation for users, it also restricts generation to chemically traceable transformations.

The results remain a proof of concept and further work will be required to apply these generative mechanistic models across engineering campaigns, for enzyme discovery, and to predict enzyme function for complex transformations. Fluxion shows that mechanism based generative methods are likely to become useful in silico tools for studying biocatalysts complementing existing retrieval based approaches.

## Acknowledgments and Declarations

## Acknowledgements

Acknowledgments

Joana Carvalho for her incredible work on the figures, and the creation of graphics to describe the model and evaluations. Esther Heid, George Bouras, Yili Shen, and Jason Nomburg for helpful discussions. AITHYRA computing which supported the development of the model through the HPC resources. AI was used for code generation, cleaning of the M-CSA data, and to facilitate with reviewing the manuscript.

## Author contributions

WJFR and ANM conceptualization, data curation, software, analysis, visualization, methodology, writing. ANM, MB, AT, JR and SL supervision, conceptualization and revisions. AT contributed to the code and model development. SHH, LH, and MM experimental set-up and experimental validation. BPF contributed to data analysis, baseline predictions and code validation. GL, SHH, and ZL contributed to the code validation and manuscript finalization.

## Data, code and materials availability

All data, code, and models will be made public on publication. If anyone would like access before publication please get in contact.

## A Methods

### A.1 Agentic data processing of the M-CSA database

Raw mechanistic data were sourced as Marvin MRV files from 734 enzyme mechanisms in the M-CSA database. Each MRV file encodes a multiple step enzyme reaction as a series of manually drawn chemical structures with annotated electron flow arrows, lone pairs, formal charges, and bond orders (including coordination bonds). We developed a custom parser that converted each step’s MRV CML into a structured plain-text format containing four sections: a list of atoms, bonds, electron flows, and metadata. Each atom record includes a local identifier, element type, 2D coordinates, formal charge, lone pair count, and radical state. A local context signature was inferred from the bond data and recorded as a compact string encoding the atom’s immediate bonding environment (e.g., C(C1,N1,O2) for a carbon bonded to carbon, nitrogen, and a carbonyl oxygen). Coordination bonds were preserved as single-bond entries marked with an identifying label. Electron flows which are encoded as directed arcs specify a source (one atom for lone pair donation, two atoms for bond cleavage) and target (two atoms for bond formation, one atom for lone pair receipt), plus a label indicating whether the transfer involved a single electron or an electron pair.

As each mechanistic step is drawn independently by human annotators, atoms that persist across steps carry no shared identifier in the raw MRV format. Establishing atom correspondence across steps by assigning each atom a stable global atom ID and a global molecule label is a prerequisite for constructing reaction representations that represent a balanced stoichiometric reaction. Atoms were labeled with a large language model (LLM), using both Google Gemini 3.1 - Pro and Google Gemini 3 - Flash on the 22nd of February, 2026. The annotator was prompted with the metadata, the plain-text representation of step *i*, and step *i* + 1, and instructed to match atoms between steps using element identity, position and context signature. The model then populates the empty global atom id and global molecule label fields in the atom data. A cascading scheme ensured consistency across the full reaction: for the first step pair, step 0 atoms were assigned their local identifiers as global IDs, and step 1 atoms were matched to them; for each subsequent pair, the annotated output of step *i* + 1 from the previous query became the fixed input for step *i* in the next query, so global IDs propagated forward without drift. Atoms genuinely new in step *i* + 1 (e.g., a water molecule entering the active site) received unique IDs of the form sNNNaXX, where NNN is the step index and XX is a unique atom identifier. After generation, structural integrity was checked by verifying that the response contained exactly two complete “ <txt>” output blocks. Second, per-step validation compared the AI output against the original input to detect lossy errors (atom or bond count changes, indicating data loss) and corruption errors (modification of protected fields such as element types or bond orders, which should never be altered). Lossy errors triggered a retry; corruption errors were resolved by force-overwriting the affected fields with the original values while retaining the AI’s global ID and label assignments. If the primary model (Google Gemini 3 - Flash) failed after an initial retry, the query was escalated to a more capable model (Google Gemini 3.1 - Pro, [40]).

The atom-mapped step files were processed by an automated consolidation pipeline to produce a unified matrix representation of each reaction. Step files were first parsed into typed data structures (Atom, Bond, ElectronFlow, StepInitial). Bond identifiers were resolved from local step-specific atom IDs to global atom IDs using the annotator’s output. Final-step states were derived programmatically by applying the electron flows to the initial bonding graph: bond orders were incremented or decremented along each flow arc, and lone pair counts were updated by the corresponding electron changes.

All unique global atom IDs appearing across any step were pooled into a single ordered atom set of size N, establishing a fixed index for every atom in the reaction. For each mechanistic step *k*, three *N ×N* matrices were constructed. The Bond-Electron (BE) matrix encodes the complete electronic state at the start of step *k*: diagonal entries hold the number of non-bonding electrons (lone pairs plus any radical electrons) on each atom, and off-diagonal entry *BE*[*i, j*] holds the number of bonding electrons shared between atoms i and j. The “delta-BE” matrix encodes the electron redistribution during step *k*, derived from the annotated electron-flow arrows. This explicit electron representation was later converted to store bond orders, as 0.5 for a radical bond, 1 for a single bond and 2 for a double bond. Lone pair donation to a bond decrements the donor’s diagonal and increments the corresponding off-diagonal pair, while bond cleavage to a lone pair does the reverse. The “final BE” matrix is the sum of the BE + delta-BE, representing the state of the electrons at the end of step *k* and the start of step *k* + 1.

Annotators often draw only the atoms directly relevant to each step, hence many atoms (cofactors, spectators, distal residues) are absent from a given step’s atom list despite being present in the reaction. Missing atoms were filled in by examining each atom’s first and last annotated appearance. If an atom had not yet appeared by step *k* it was represented using its bonding state at its first appearance (the pre-reaction state); if it had already disappeared before step *k* it was represented using its bonding state at its last appearance (the post-reaction state). This produces BE matrices in which every atom is present at every step, making each step a complete balanced snapshot of the entire reaction rather than a partial view. Coordination bonds (dative bonds from a Lewis base donor to a metal acceptor) were handled separately from covalent bonds. Because the Marvin software annotates lone pair counts to include the electrons involved in a coordination bond, adding those electrons again to the donor atom’s diagonal would double-count them and produce incorrect formal charges. Coordination bond electrons were therefore excluded from the BE matrix entirely. Instead, coordination bond topology was recorded in a binary COORD matrix (1 where a coordination bond exists between atoms i and j, 0 elsewhere), which is stored alongside the BE matrices and can be used for downstream analyses requiring metal-ligand connectivity, but was not used in model training.

Ideally the final BE matrix of step *k* equals the BE matrix of step *k* + 1 for all shared atoms, reflecting a consistent electron count across the reaction sequence. Discontinuities that typically arise from annotation ambiguities or steps in which the annotator chose a different resonance structure were detected by comparing final *BE*[*k*] with *BE*[*k* + 1] for each entry. When a discontinuity was found, both candidate matrices were scored for chemical plausibility (penalizing hydrogen valence violations, exceeded maximum bonding electrons, and negative entries), and the lower-violation representation was propagated to replace the higher-violation one. This process was applied iteratively until no discontinuities remained.

Consolidated matrices were checked against a set of chemical constraints: bonding electron counts must not exceed the element’s maximum valence; lone pair counts must be non-negative; and electron conservation must hold across each step (total electrons in final *BE*[*k*] must equal total electrons in *BE*[*k* + 1] for atoms present in both). Reactions failing validation were flagged and excluded from downstream dataset generation. We acknowledge that this automated pipeline does not guarantee full chemical correctness, and likely contains noise in the form of incorrect protonation states. We recognize the contribution of Hartley et al. [20] in manually correcting inconsistencies across the database, and envision the dataset quality could be improved by integrating their corrections.

Each mechanistic step of a consolidated reaction yields one training pair. For step *k*, the reactant state **x**_0_ is the BE matrix at the start of the step and the product state **x**_1_ is the corresponding final BE matrix, **x**_0_ + Δ**BE***_k_*. Because consolidation pools every atom of the reaction into a single indexed set, both matrices span the whole mechanism and the pair is atom-aligned by construction, with no atom mapping required at training time. Hydrogens were made explicit before pairs were formed. For every C, N, O, S, P, B and Se atom the suppressed hydrogen count was recovered from the valence identity above, evaluated on the first state of the mechanism, and that many hydrogens appended to the pooled atom set with a single bond to their parent atom. Hydrogens already drawn by the curators are left untouched, as are metals and bare protons. The same hydrogen bonds are written into every state of the trajectory rather than recomputed per step. Every step’s Δ**BE** is therefore unchanged by the reconstruction, and only the constant valence context around it differs. For each reaction, an additional terminal step, which describes no change to the reaction’s product BE matrix is added to the pool of data, so that the model is trained to be able to predict when a reaction ends. The final dataset contains a total of 2,747 steps, across 599 reactions.

**Supp. Table 1:** Summary of M-CSA mechanistic entries processed by the agentic reconstruction workflow.

| Status | Count | Percentage (%) |
| --- | --- | --- |
| Reconstructed and retained | 599 | 81.6 |
| Without warnings | 497 | 67.7 |
| With $\geq 1$ warning-flagged step | 102 | 13.9 |
| Excluded — failed chemical validation | 133 | 18.1 |
| Excluded — no parsable mechanism input | 2 | 0.3 |
| Total M-CSA mechanistic entries | 734 | 100.0 |

**Supp. Table 2:** Breakdown of errors observed among failed M-CSA entries. Error categories are not mutually exclusive, and individual entries may contain multiple errors.

| Error | Count |
| --- | --- |
| Valence violation (too many bonds) | 83 |
| Conservation error (electrons not conserved) | 68 |
| Total electron mismatch across reaction | 68 |
| Hydrogen bond violation | 30 |
| Negative bond electrons | 25 |
| Exception during processing | 4 |

### A.2 USPTO data processing

Pretraining used the atom-mapped USPTO mechanistic dataset curated by Joung et al. for FlowER, with their published train, validation and test partition adopted unchanged [15]. No reactions were added, removed or re-assigned. Each mapped step was re-expressed in the bond-electron representation described above: the reactant and product SMILES were Kekulized, off-diagonal entries set to the bond order between each atom pair, and diagonal entries set to the non-bonding electron count implied by the valence equation. The curation draws hydrogens explicitly, so no hydrogen reconstruction was required and the pretrained encoder sees the same explicit-hydrogen valence context as the M-CSA fine-tuning data.

### A.3 EnzymeMap data processing

Additional atom mapped enzymatic reaction data were derived from EnzymeMap v2 [41] to train the reactant to product single step reaction model. Atom mapped reactions labeled as directly reversed and reactions without an associated UniProt protein sequence were excluded. This yielded a total of 7,825 protein sequences, mapping to 12,632 distinct reaction SMILES, of which there are 38,187 protein-reaction associated pairs. A total of 51,610 records were used in practice, as records were not de-duplicated by reaction or protein-reaction pair. Catalytic residues for each sequence were taken from a combination of Squidly predictions [42], UniProt catalytic residues and substrate binding annotations. UniProt binding-site annotations were used to identify putative cofactors. Records were then constructed in the BE matrix format described above for the USPTO, and M-CSA datasets. Each resulting EnzymeMap record contains only the reactant and product endpoint rather than an experimentally curated sequence of elementary steps. Added catalytic residues and cofactors were included identically at both endpoints and therefore act as spectator features. Finally, these were subset into train and test splits using the definitions in the CARE benchmark, taking the easy split from task 1, resulting in 28,837 training, 1,601 validation, 1,596 test enzyme-reaction pairs [10].

### A.4 Esterase data processing

Code and data were downloaded from https://github.com/samgoldman97/enzyme-datasets, based on the papers [34] which used raw data from Martínez-Martínez et al. [32]. We used the raw data from Goldman et al. [34] and reprocessed it to include normalized activity data. Sequences greater than 1000AAs were dropped (N=1). Catalytic sites were annotated using Squidly and BLAST ensemble, using a threshold of 0.3 for BLAST. Non-zero activity values were normalized by performing a Z transform, clipping values greater than 3SD above the mean, these were then min-max transformed. As no products exist by default in the dataset, but given the esterase reaction is the breaking of an ester bond, we opted to generate the products for those where this was possible. To balance the reaction (ester to acid and alcohol) we add water to the LHS of the reaction (O, SMILES fills valence with implicit hydrogens using Chem.MolToSmiles(Chem.AddHs(Chem.MolFromSmiles(“O”)))). The reactions are then checked for being balanced by using rdkit. This is run for each of the training-test-validation splits. When the data are used for finetuning, no imputed negatives are introduced as in the training from EnzymeMap as there are true negatives in the dataset. The activity head has a mask such that it is only learnt when the label is 1 to avoid collapse to 0 owing to imbalanced datasets. All evals are performed with 64 samples except where otherwise stated as in the esterase and phosphatase evaluation (Fig. 4).

### A.5 Phosphatase data processing

Phosphatase data is from Huang et al. [33], but the processed data were sourced from Goldman et al. [34], as above. The processing for the Phosphatase dataset is the same as the esterase dataset except using OP(=O)(O)O to balance the reaction as a co-product. The atom balancing for the phosphatases is: R-OPO_3_ + O → R-OH + OP(=O)(O)O. Training is performed as above just with a different dataset.

### A.6 EnzEngDB data processing

EnzEngDB [39] was downloaded from https://doi.org/10.5281/zenodo.17310823. Cofactors are not by default in here, so we used a combination of manual and automated approaches to add the cofactor to each row in the EnzEngDB. Reactions were filtered for being atom-mapped and balanced, removing unbalanced rows. We created two tests for the EnzEngDB, first seeing if there is a difference in the predicted reactivity between the start and end of a campaign, selecting the main reaction and the one with the least mutations vs the most mutations. For EnzEngDB no catalytic residues were assigned rather the mutations from the campaign were used for the PLM conditioning (but not added as spectator atoms in the matrix).

### A.7 Data processing from NHC-Mediated Radical Acylation Catalyzed by Thiamine- and Flavin-Dependent Enzymes

Uniprot IDs were taken from Table S1. Screening results of ThDP/FAD-dependent enzymes [37]. The sequences were then taken from Uniprot and used with the reaction as defined in the EnzEngDB for screening. No catalytic residues were defined for these sequences. Yield was reported as in the figure and considered as active if detected.

**Supp. Table 3:** Results from the SI of [37] alongside whether Fluxion generated a hit for this reaction, and at what rank the hit was observed.

| id | hit | rank | fitness |
| --- | --- | --- | --- |
| C0ZWJ1 | TRUE | 3 | 0 |
| P0CH62 | FALSE | NaN | 0 |
| D4YIP5 | TRUE | 4 | 0 |
| P07003 | TRUE | 3 | 0 |
| C1A0V1 | FALSE | NaN | 0 |
| Q88F03 | FALSE | NaN | 0 |
| P0AEP7 | TRUE | 2 | 0.12 |
| P08142 | TRUE | 2 | 0.19 |
| P0DP90 | TRUE | 2 | 0.2 |
| A0A067Z243 | TRUE | 4 | 0 |
| Q5SJ01 | TRUE | 4 | 0.2 |
| Q47SB8 | TRUE | 4 | 1.3 |
| A0A919QSP1 | TRUE | 4 | 0.88 |
| D6Y6A4 | TRUE | 3 | 2.4 |

### A.8 Train-test-validation splits from EzSpecificity

To enable comparison to existing methods, we downloaded splits from EzSpecificity [4] for the esterase and the phosphatase datasets, these splits used IDs from each respective dataset for splitting, this was downloaded from: https://drive.google.com/drive/folders/1-lfpKPhmf_P837UBfgUt_7R00_f45DtA, June 2026.

### A.9 Benchmark curation for M-CSA

To assess model performance, an in silico benchmark was created with three unique train, validation and test splits designed to span varying levels of difficulty in enzyme mechanism prediction based on the pairwise mechanism similarity defined by Ribeiro et al. [19]. Similarity is defined as the intersection-over-union, or Jaccard similarity, between the unordered sets of mechanistic “arrow environments” associated with each reaction, irrespective of their ordering across mechanistic steps. These pairwise similarities are provided as an edge list, where each edge connects two M-CSA entries with non-zero mechanistic similarity. The “easy” test set was derived by sampling reactions with greater than 0.5 and less than 0.9 similarity to other reactions in the dataset, with fewer than 100 total heavy atoms and no more than 10 mechanistic steps. The “medium” test set sampled reactions with greater than 0 and less than 0.5 similarity and was restricted to mechanisms containing more than four steps. The “hard” test set was drawn primarily from reactions with no mechanistic-similarity edge to another M-CSA entry, together with ten mechanisms selected for extreme atom or step counts. Because each similarity pair is listed only once, with one mechanism arbitrarily designated as the source, our eligibility check, which scanned source-side entries only, missed similarity edges for 65 of the 599 reconstructed reactions that appeared exclusively as edge targets. These reactions were therefore treated as having unmeasured similarity during split construction. As a result, 21 of the 80 hard-test mechanisms retain a partial mechanistic analogue in the training or validation set (mean maximum similarity 0.43 among these 21, including two identical mechanisms), while the remaining 59 have zero measured similarity. The hard set therefore has a mean maximum similarity to training of 0.10 rather than zero (Fig. 2C). The easy and medium splits are defined by the presence of similarity. They are therefore not compromised by this omission.

Each test set was fixed at *n* = 80 reactions and assembled in three stages. First, minimum allocations were set by hand for each top-level EC class to guarantee a minimum level of class representation, rather than allowing the sample to follow the pool’s skewed composition, where a class had fewer eligible reactions than its allocation, all available reactions were taken (Supp. Table 4). Second, reactions were drawn at random within each class up to its allocation. Third, the set was topped up by random draws from the remaining eligible reactions until 80 were selected. For the hard set, ten reactions representing the extremes of the dataset in total heavy-atom count or number of mechanistic steps were added before quota sampling and count towards the 80. All draws used a fixed random seed. The remaining reactions formed the training pool, from which 10% were sampled at random as a validation set; the validation split is therefore not EC-balanced. This gives 468 training, 51 validation and 80 test reactions for the easy, medium and hard splits.

**Supp. Table 4:** Minimum per-class allocations for each test set, with the random top-up to *n* = 80.

| EC | Class | Easy | Medium | Hard |
| --- | --- | --- | --- | --- |
| 1 | Oxidoreductases | 5 | 15 | 15 |
| 2 | Transferases | 10 | 8 | 10 |
| 3 | Hydrolases | 20 | 30 | 10 |
| 4 | Lyases | 5 | 10 | 10 |
| 5 | Isomerases | 5 | 8 | 10 |
| 6 | Ligases | 0 | 3 | 2 |
| 7 | Translocases | 1 | 0 | 2 |
| Allocated |  | 46 | 74 | 59 |
| Random top-up |  | 34 | 6 | 21 |
| Total |  | 80 | 80 | 80 |

### A.10 Bond-electron matrix representation

Here we define the final bond-electron (BE) matrix representation used in training. A reaction state over *N* atoms is represented by a symmetric BE matrix:

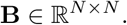

The diagonal entry *B_ii_*is the raw number of non-bonding electrons on atom *i*, including lone pair and unpaired radical electrons. The original representation used in the agentic cleaning pipeline for off-diagonal bonds is halved in order to represent bonding electrons as bond orders, and thus the off diagonal entry *B_ij_* is the bond order of the bond between atoms *i j*. Closed-shell single, double, and triple bonds are represented by bond orders 1, 2, and 3, respectively, whereas a one-electron bond arising from radical or homolytic chemistry is represented by a bond order of 0.5. The total number of valence electrons encoded by **B** is:

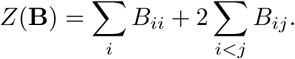

Equivalently, *Z*(**B**) is the plain sum of all entries in the full symmetric matrix. Symmetry causes each bond entry to be counted twice, such that a bond of order *B_ij_* contributes 2*B_ij_* shared electrons.

A reaction step from **x**_0_ to **x**_1_ is electron-conserving when

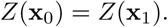

A half-bond with *B_ij_*= 0.5 contributes exactly one electron. Therefore, this electron-counting convention applies identically to closed-shell and half-integer radical representations.

In the Marvin-derived M-CSA mechanisms, a hydrogen atom is drawn explicitly only when it participates directly in the mechanism. Spectator C–H bonds are therefore generally absent. For each atom *i*, the implicit hydrogen count is reconstructed using the valence identity

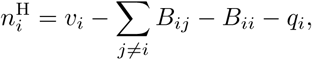

where *v_i_*is the number of valence electrons associated with the atom and *q_i_* is its formal charge. The reconstructed hydrogen atoms are appended to the matrix as single bonds, and are added into every state of the corresponding mechanism.

### A.11 BE to BE Schrödinger Bridge

We model an elementary reaction step as a stochastic transport in BE-matrix space that begins at the reactant. Reactants and products of a step share a pooled atom index, so the two states differ only by a sparse redistribution of electrons, and a single reactant can admit several chemically reasonable outcomes as a distribution of plausible products. Therefore, we learn the transport as a stochastic diffusion bridge between the paired endpoints following the Image-to-Image Schrödinger Bridge of Liu et al. [18]. Closely related work by Somnath et al. [43] independently developed diffusion Schrödinger bridges for aligned endpoint data. The network is conditioned on the current state and time, and optionally augmented with protein language model embeddings.

During training a time point is sampled uniformly on an interval between a small *t*_min_ and 1 − *t*_min_ which avoids numerical issues at around *t* = 0 and *t* = 1.

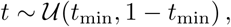

and the bridge state at that time is

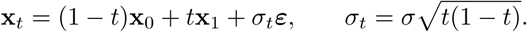

Here *ε* is Gaussian noise restricted to the valid matrix entries m, centred over them so that it carries no electrons, *Z*(*ε*) = 0, and symmetrized,

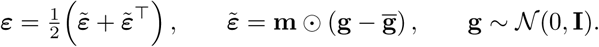

The network regresses the endpoint displacement rather than the bridge noise,

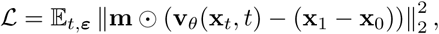

summed over matrix entries and averaged over the batch. This departs from Liu et al. [18], where the network predicts *ε*. Predicting x_1_ x_0_ means the zero-sum symmetric projection applied by the output head acts directly on the supervised quantity. Reported models use *σ* = 0.3 and *t_ε_* = 0.005.

At inference, time is discretised into *S* steps over

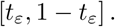

At each time *t*, the model estimates the clean product matrix as

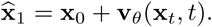

For a transition from *t* to *t*^′^ = *t* + Δ*t*, the posterior mean is

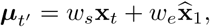

where

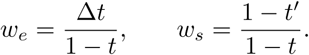

The next bridge state is sampled as

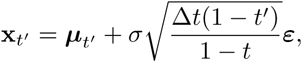

where *ε* is again symmetric, zero-centred, and electron-conserving.

During inference, generated products are rounded to a discrete electron conserving state, as per FlowER’s implementation, except to allow radical chemistry, the upper triangle, off-diagonal entries are rounded to the nearest half, which preserves a one-electron bond as a bond order of 0.5. The diagonal is rounded to the nearest integer, and all entries are clamped at zero. Since rounding generally disturbs the target electron count taken from the reactant, *Z*(x_0_), the residual is absorbed by the diagonal alone, incrementing or decrementing entries in order of the fraction discarded and skipping any that would be negative. If possible, each generated matrix is converted to a canonical-SMILES representation before ranking using RDKit, and matrices mapping to the same SMILES have their counts pooled before ranking [44].

Each reactant BE matrix is sampled *R* times. Every run begins at the same x_0_ and integrates the same learned field, so proposal diversity is produced entirely by the noise injected at each posterior step rather than by any variation in the starting point. The *R* decoded matrices are then ranked by the number of draws that produced them. The resulting count is the model’s implied frequency for that outcome and supplies the ordering used by all top-*k* reporting, so top-*k* accuracy is bounded above by the number of distinct candidates a reactant receives and cannot exceed *R*. All reported results use *R* = 64 draws and *S* = 50 bridge steps.

### A.12 Model architecture

The model uses a GraphGPS-inspired [45] graph Transformer that uses a sequential combination of a edge-aware message-passing, operating on continuous-valued hidden representations, and global self-attention over valid atoms followed by a feed-forward network (Fig. 1C). The model represents the full BE matrix as a graph and the static reactant graph is constructed from atom pairs satisfying

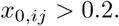

This threshold ensures that half-bonds with bond order 0.5 are included as graph edges for radical reactions. The graph connectivity is precomputed before noise is injected in to the matrix, so whilst noise does affect edge and node features, it does not lead to dynamic changes in the graph connectivity. Atom elements are encoded with an embedding that is learnt across the pretraining with the USPTO dataset, and the electron/bond values in the diagonal and off-diagonal matrix entries are expanded using Gaussian radial basis functions. Between the GPS layers an optional cross attention layer is inserted which integrates the model’s graph representation with precomputed ESM2 [22] per-residue token embeddings and mean-pooled sequence embeddings (model name “esm2-t36-3B-UR50D”, embedding dimension = 2560). The model can be configured to accept a variable number of tokens enabling other representations to be added in the future. The final benchmark Fluxion models all had 6 GraphGPS-style layers, whilst PLM models had cross attention heads after layers 2, 4 and 6.

The final atom representations are decoded into a dense matrix-valued electron-displacement prediction using separate constructions for diagonal and off-diagonal entries. Let h*_i_* R*^d^* be the final representation of atom *i*. For every ordered atom pair, a preliminary pairwise output is computed as

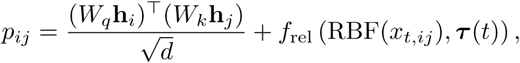

where *τ* (*t*) is the sinusoidal time embedding and *f*_rel_ is a multilayer perceptron. This dense pairwise head is evaluated for all valid atom pairs, not only edges in the static message-passing graph, allowing the model to predict both the breaking of existing bonds and the formation of new bonds.

Diagonal entries, which describe changes in non-bonding electron counts, are generated using a separate attention operation. For each atom,

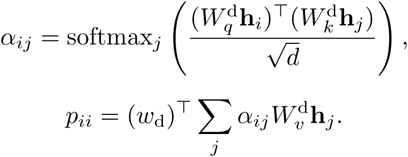

Thus, the predicted change in the non-bonding electrons of an atom may depend on the representations of every other atom in the reaction. These values replace the diagonal entries produced by the pairwise branch.

All reported models share one encoder: 6 pre-norm transformer layers with a graph message-passing branch, hidden width 256, 16 attention heads, a feed-forward width of 1024, and a 64-dimensional sinusoidal time embedding added to the atom states. Where protein language model conditioning is used, ESM2-3B residue embeddings of dimension 2560 enter by 8-head cross-attention re-injected every two encoder layers.

### A.13 Protein language model conditioning

Protein language model embeddings of each enzyme’s sequence are used as optional conditioning, to enable the enzymatic context of the sequence to guide the conditional flow. Catalytic-residue assignments were compiled from the M-CSA records for all entries, recording each residue’s position against its own UniProt accession so that enzymes with catalytic residues on several subunits are represented in full. Reference sequences for those accessions were retrieved from UniProt. ESM2 (esm2_t36_3B_UR50D, embedding dimension 2560) was then run once per sequence, and the per-residue token at each catalytic position was retained together with the whole-sequence mean-pooled embedding. This yields one native embedding set per reaction and subunit.

Training reactions may additionally be conditioned on homologous enzymes to the M-CSA canonical enzyme, to assess if the augmented increase in variance of enzyme sequence conditioning would improve generalization. The 599 reference sequences were searched against UniRef90 [46] with BLASTP (E-value 10^−3^, up to 2000 targets per query) [47], retaining hits with at least 30% identity and 50% query coverage. For each reaction and subunit the hits were binned by identity to their own query into 30–50%, 50–70% and 70–90% intervals and up to 50 were kept per bin, preferring homologues with an annotated RHEA reaction and, among those, experimental over automated evidence. Catalytic positions were transferred from the query to each retained homologue through a MAFFT pairwise alignment [48], and ESM2 was run on the homologue sequence to give a further embedding set with the same structure as the native one. Homologues augment training reactions only whilst validation and test reactions are always conditioned on the native embedding of their own reference enzyme, never on a homologue. To prevent leakage, a homologue is admitted only if it shares no more than 0.3 identity with the reference sequence of any validation or test reaction. Because the easy, medium and hard splits partition the same 599 reactions differently, the second constraint is evaluated per split. At training time, a reaction’s admissible sets are grouped by subunit and one set is sampled per subunit for each example, so that multi-subunit enzymes are seen as combinations of subunit sequences. The resulting tokens, one per catalytic residue plus the mean-pool token, are projected to the encoder width and read by cross-attention modules inserted after every second encoder layer, each atom attending to all tokens.

### A.14 Multi-step mechanism inference

To enable the generation of full reaction mechanisms, we search over iterated single-step predictions. Beginning from the reactant, the beam search builds a directed graph of states. At each depth, every state held for expansion is passed to the single-step model sampler. The model draws a specific number of proposals, and returns them pooled by identity and sorted by how many draws produced each unique proposal. Thus each distinct outcome has a probability used to rank them. Then the subsequent best candidates are used to sample the next step, controlled by the depth parameter. A path is complete when a terminal step is reached. Terminal steps are steps where no change occurs, however other nodes are still searched when a terminal node is reached and most probable. If the proposal lands on a state already seen in the graph, the edge is recorded but the state is not queued for further sampling. All resulting pathways in the search are then evaluated by their mean per-edge log-probability of each step in the pathway. Optionally, pathways can be evaluated by filtering only those which reached a known product, to allow for mechanism prediction for a known reaction outcome.

### A.15 Start-end inference and training setup

In comparison to the mechanism problem, we also sought to test training and evaluation on going directly from the start state to the final state, here the catalytic residues are only spectators as these should not be changed at the final state (in comparison to the mechanism formulation where they can be involved in the mechanism). Training was performed across USPTO (N=237,207 for training), and then using this to warm start a training on EnzymeMap (N=28,837 reaction-enzyme pairs for training, and 1,601 for validation and 1,596 test), training each until a maximum of 800 epochs. USPTO was warm started from the output of the enzyme mechanism training (i2sb_6L_256d_scratch_sigma03), and saturated at 0.889 at epoch 550, while training for EnzymeMap reached validation loss of 0.666 at epoch 480. The best checkpoint was determined using the metric val_blended_top1_step_acc which evaluates the generated SMILES vs the true SMILES. I^2^SB was used with ESM2 model esm2_t36_3B_UR50D for the PLM and the catalytic residues. Learning rate of 5 × 10^−5^ is used with EMA for USPTO, and 2 × 10^−5^ for EnzymeMap.

### A.16 Evaluation metrics

When evaluating model performance, the discretization of the model output allows direct comparison to the reference product for the ground truth reaction steps. When evaluating stepwise performance, the generated BE matrix is converted to a canonical-SMILES representation and chemically infeasible BE matrices are discarded after failed conversion to SMILES. The resulting generations are scored by their accuracy in generating the correct reference product, in the top-*k* most sampled outcomes. Not every ground truth state can be converted to a SMILES, and reconstruction also fails for some hypervalent or charged intermediates. Discarding these would score the model on an easier subset than it was trained on, so we report a top-*k* that scores each step by SMILES where the reference state can be reconstructed into SMILES and by exact BE match where it does not. We ablate the performance increase from the SMILES conversion in our ablations.

For full reaction mechanism prediction using the beam search, we report whether the reference path appears among the top-*k* ranked paths for all paths where the known product terminal is reached. The top-*k* accuracies for a reaction is zero by default if the product is never reached by the model. Additionally, we report whether the reference product appears in any pathway among the top-*k* ranked paths among all generated paths.

### A.17 Mechanism prediction baselines and ablations

Baselines and ablations were established to isolate the contributions of the training objective, model pretraining on the USPTO, and the sampler’s noise scale. First we replaced the stochastic bridge with a deterministic model: the same encoder, output head and conservation projection were used and initialized from the same USPTO checkpoint, but the model makes a single forward pass on the clean reactant and regresses the whole-step displacement directly, so its prediction is x_0_ + *f* (x_0_). Because it is deterministic, repeated draws are identical and it returns exactly one candidate per step, so its top-*k* accuracy equals its top-1 for every *k*. Secondly, we ablated the USPTO pretraining and trained directly on the M-CSA from random initialization. The learning rate, warm-up length and gradient clipping necessarily differ from the fine-tuning schedule, which does not train a model from scratch. Thirdly, we varied the noise injected by the posterior sampler while holding the trained (native PLM) model fixed: the calibrated noise of the *σ* = 0.3 model is multiplied by a factor between 0.5 and 2.0, giving effective values from 0.15 to 0.6.

In order to compare our overall architecture to FlowER’s, we attempted to fine tuned the most recent publicly available checkpoint of the FlowER model, released in May 2026. First the easy data split was converted to FlowER supported SMILES. 20.7% of steps were dropped by FlowER’s data preprocessing pipeline, for having illegible SMILES caused by hypervalence or half bonds, or chemical inconsistencies that may be introduced by the agentic data cleaner. Attempts to fine tune the model collapsed to predicting no change for each step in training, even when applying a weight to the FlowER’s MSE loss for changed cells in the matrix displacement.

To evaluate the baseline performance of a rule-based mechanism prediction tool such as MechFind, training specific rulesets were generated independently for the Easy, Medium, and Hard splits. For each split, rules were generated exclusively from the M-CSA entries in the training set. Validation and test entries did not contribute rules. Steps were generated with a constitutional radius of 1 and both the forward and reverse steps were included together with the predefined protonation and deprotonation rules used by MechFind. Duplicate rule vectors were collapsed into a single rule. Test-set mechanism steps were then converted using the same constitutional radius of 1. We then calculated the overall step coverage and mechanism coverage for each test set as the proportion of total steps which have an exact matching rule, and the proportion of total mechanisms that have all rules with a matching step. Because the stepwise benchmark includes the terminal reaction step used by Fluxion to determine reaction termination at inference, the numerator and denominator of the fraction is increased by 80, to ensure the comparison is fair. These values represent optimistic rule-availability ceilings as they assume perfect selection and ordering of the required rules and do not measure whether the inference procedure actually discovers the annotated mechanism. This ceiling is defined within MechFind’s constitutional radius-1 representation. It does not evaluate stereochemical outcomes or plausible mechanisms that are not among the mechanisms annotated in M-CSA.

### A.18 CYP regioselectivity benchmark

A new benchmark for regioselectivity in CYP enzymes was derived from the substrate/reaction promiscuity benchmark designed by Mahood et al. [9]. The source set comprises 2921 atom-mapped enzyme reaction pairs spanning 506 distinct substrates and 768 UniProt entries. From these we retained reactions representing a single, unambiguous aliphatic C-H hydroxylation: an atom-mapped, non-aromatic carbon that gains exactly one oxygen neighbour in the product. Reactions with multiple such carbons, i.e. diols, multi-site oxidations, and aromatic hydroxylations were excluded as the target carbon cannot be identified without ambiguity. This yields 1045 training, 145 validation and 173 test enzyme-reaction pairs. With this data, we formulate the task as so: For each substrate and enzyme pair in a hold out test set, the model ranks every candidate position on that substrate and is scored on where the enzyme’s own observed site(s) fall in the ranking. Candidate sets range from 5 to 11 possible positions, depending on the held out substrate. Thus the task is set up to assess the utility of a model which can report probability distributions over multiple product outcomes.

To measure generalization to unseen chemistry, we held out substrates such that for a held out molecule, no training substrate or reaction exists. To select the substrates for testing, substrates were ranked by the number of scoreable hydroxylation positions each would contribute. Five substrates, including arachidonate, cholesterol, *β*-amyrin, progesterone and linoleate, were selected to maximize positional diversity and relative chemotype spread (see 3C and 3D). Near relatives to these substrates were removed from the training data by first filtering out any substrates with greater than 0.85 tanimoto similarity, and further reduced by removing substrates with a maximum common substructure (MCS) overlap of greater than 0.90. Maximum common substructure was determined as the jaccard index over heavy atoms in the maximum common substructure of two molecules. This filter removed 66 similar substrates (444 reactions) from the data. Additionally, hold out test enzyme reaction pairs were redundancy reduced to less than 95% pairwise sequence identity, effectively reducing near duplicate enzymes so the result isn’t dominated by clusters of near identical sequences. The final test set comprises 144 enzymes, with 173 unique reactions. Enzymes are deliberately not held out and the holdout is over substrates, so an enzyme may appear in both splits acting on different molecules.

### A.19 CYP regioselectivity inference and evaluation

Each reaction was represented as a single elementary mechanism step. The modeled transition is the hydrogen atom transfer (HAT) in which the ferryl oxygen abstracts a hydrogen from the target carbon. This is the step which determines the regioselectivity. A BE matrix was constructed by transplanting the substrate onto a constant P450-BM3 Compound-I heme scaffold extracted from M-CSA entry 699 [26]. Fluxion’s bridge model was then sampled 2048 times per case for the reactant under each enzyme conditioning, and draws were binned by which carbon lost a hydrogen, not by the exact product matrix. The primary metric is the tie-aware set AUC: over all (correct, incorrect) candidate-position pairs for a given enzyme, the fraction in which the correct position scores higher, with ties counted as one half. This is the normalised Mann-Whitney U statistic. We note that the transplanted reactant offers many more abstract-able aliphatic C-H carbons however, the candidate set is the set of reported outcomes, not the set of chemically accessible ones.

Fluxion was compared with three baselines: random ranking, a substrate-transfer baseline, and a hybrid sequence/substrate-transfer baseline. The substrate-transfer baseline is the chemical analogue of a homology baseline, ignoring the enzyme entirely: for a held-out substrate, the most similar training substrates are identified by MCS, and their per-atom site distributions are mapped onto the held-out molecule through the atom correspondence. The hybrid baseline extends this to use the query enzyme as well, scoring each training enzyme-substrate pair by the product of its sequence similarity to the query enzyme and its substrate similarity to the query substrate, and transferring observed positions from the highest-scoring pairs by the same correspondence. Both transfer baselines are reported in two variants, transfer from the single closest donor and a similarity-weighted pool over the ten closest, which differ substantially because 83% of training substrates have only one recorded hydroxylation site, so a single donor can score at most one candidate and leaves the remainder tied. In both cases positions are transferred through two alignment frames combined by element-wise maximum: the MCS atom correspondence, and, for acyclic substrates, a w-frame indexing each carbon by its topological distance from the terminal methyl. Finally, we report a transductive ceiling: for each test enzyme, the popularity of each position across the remaining test enzymes acting on the same substrate. This measures the prior distribution of reactivity over substrate positions within the test population and is reported as a reference value rather than a competing baseline.

### A.20 CYP regioselectivity sequence conditioning variants

Sequence embedding conditioning was constructed at varying levels of granularity in order to assess the effect of adding additional embedding features from sequence positions known to be associated with substrate recognition in CYPs. Specifically, three levels of sequence conditioning were tested: The “substrate recognition site” (SRS), “pocket” and “catalytic” levels. The SRS positions include sequence positions across 6 canonical regions of the CYP fold, which are recognised to be involved in substrate recognition of CYPs. They were assigned using a modified version of the BoltzCYP sequence alignment workflow described by Mahood et al. [9] Reference sequences, SRS anchor subsequences and precomputed kingdom-specific MAFFT alignments were adopted from the accompanying BoltzCYP implementation. We transferred the SRS positions to the full set of CYP sequences in the original dataset by selecting the closest reference sequence for each enzyme and projecting its padded SRS alignment onto the ungapped target sequence. The ungapped positions were used to generate the set of ESM2 tokens used for conditioning these reactions. The pocket conditioning was constructed as a reduced form of the full SRS conditioning, including only four of the 6 SRS regions: “SRS1”, “SRS4”, “SRS5”, and “SRS6”, together with the conserved catalytic glutamate, arginine and heme-coordinating cysteine. The catalytic level included the positions of the conserved catalytic glutamate, arginine and heme-coordinating cysteine. All levels of conditioning included an additional mean pooled embedding derived from the full set of SRS tokens. Training and testing was done independently for each level of conditioning and their accuracy in predicting regioselectivity of held out CYPs were reported separately. We acknowledge that this homology transfer is likely to have included incorrectly identified SRS positions for diverse CYPs, but assume that these embeddings might still contain learnable signal.

### A.21 Finetuning on the esterase and phosphatase datasets

See above for processing and sourcing of each dataset. Each of the folds were evaluated training on only the training set split from the dataset from either the PLM aware (model trained with USPTO and EnzymeMap) or just the USPTO version. The following parameters were passed to finetuning; here the heads were additionally trained. AdamW, *β*_1_ = 0.9, *β*_2_ = 0.998, weight decay 0.01, gradient clipping 1.0, batch size=32, bfloat16 mixed precision, EMA decay 0.999), with the head parameters in their own optimiser group at a 10× learning-rate multiplier (LR of 5e-5). The reactivity and activity losses were weighted at 0.5 each against the flow objective. The enzyme head had a dimension of 8, a LayerNorm-bottlenecked projection of the ESM-2 mean-pooled sequence embedding, supervised by an auxiliary term (weight 0.5) on the enzyme’s train-fold promiscuity and added zero-initialised to both readouts. An interaction cross attention layer is between the PLM and the learnt representation prior to the BE matrix, in which the attention query is made substrate-dependent and augmented with an explicit rank-1 bilinear cross-term between the pooled substrate representation and the attended enzyme context. Checkpoints were selected on the validation head score, evaluated every five epochs, and the selected model was scored with eight sampled trajectories per record.

### A.22 Evaluation and baselines on the esterase and phosphatase datasets

Two baselines were calculated for the enzyme datasets. First the base rate is calculated for the AUPR score, namely the positive rate within the train set. Next, baselines are calculated for each test set based on the training set. Namely the enzyme rate, i.e. for the training set, how many substrates was this enzyme active on, similarly, for the held out enzymes, the baseline rate is calculated as the substrates mean rate of activity/reactivity from the train set. For the random set neither are held out, here the better of the two is taken. Esterase size 146 (enzymes) × 96 (substrates) = 14,016, of these only 93 substrates were able to be atom balanced resulting in 13,578 atom-balanced samples, where 22.4% are active; for the phosphatase there are 218 (enzymes) × 165 (substrates) = 35,970, which drops to 20,056 when passing through the atom balancing filter (92 substrates passed), with approx. 16.5% active. The atom-balance filter removes approx. 3% of the esterase grid and approx. 44% of the phosphatase grid. Notably, post sample filters are uneven across splits.

Next we opted to test whether the embeddings from Fluxion were informative for predicting reactivity. Here we used either the embeddings pooled from the output (atom pooled encoder output), the embeddings from the reactivity head (atom pooled encoder output though layer norm, and cross attention context, and mean pool from the ESM embeddings, esm2_t36_3B_UR50D with 2560 dimensions). This was passed to a gradient boosting model from sklearn (HistGradientBoostingClassifier) using 4 threads. The baseline for this concatenates ESM2 embeddings and Morgan fingerprints, or Unimol2[49] embeddings which are passed to a GBM model with consistent architecture. The model’s parameters were held consistent across all runs, max_iter=50, learning_rate=0.06, early_stopping=True, validation_fraction=0.15, random_state=0.

---

**Figure S1:**
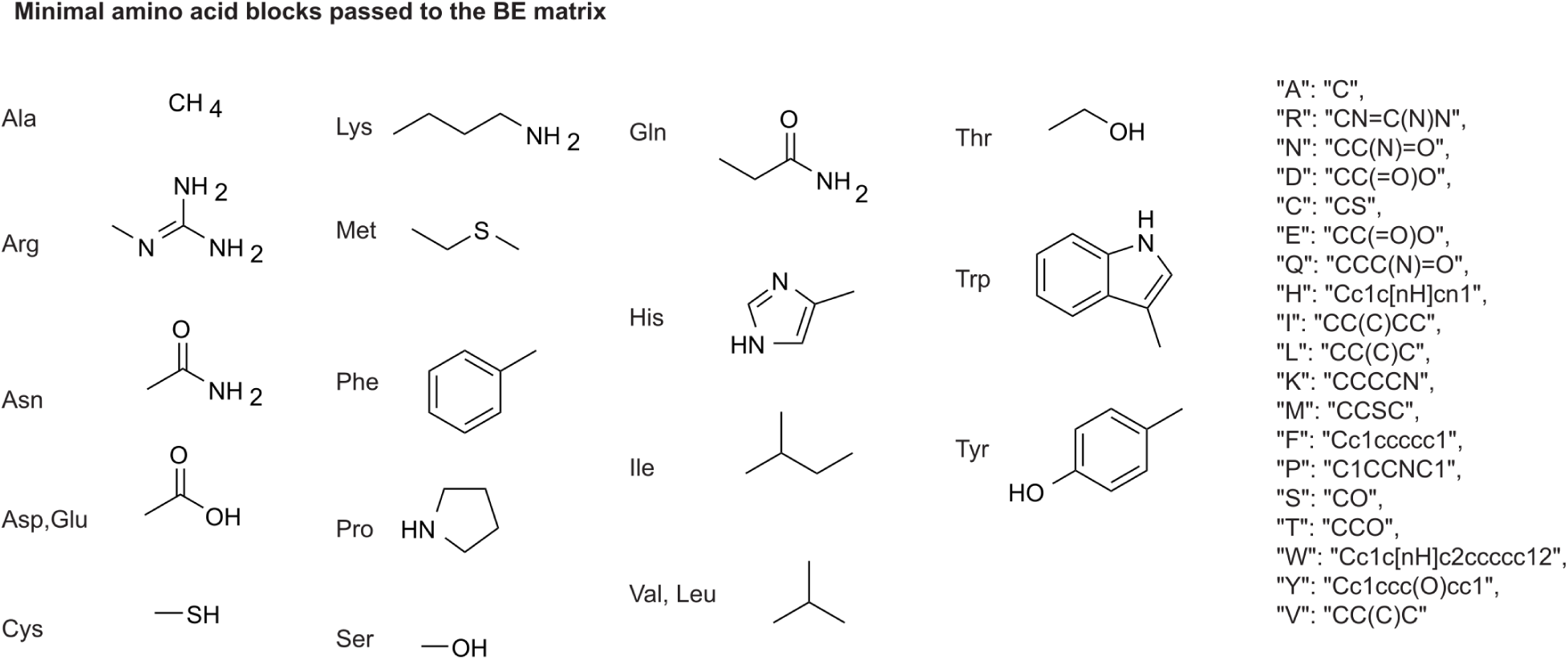
Additional details of the Fluxion representation and training framework. **A**) Amino acid SMILES representation used to construct the bond-electron (BE) matrix inputs to Fluxion. **B**) The training and experimental design to validate the Fluxion framework shown alongside the T-SNE compressed latent space learnt across the three datasets. **C**) Illustrative mechanistic step from M-CSA entry 1. M-CSA mechanistic annotations were processed using an agentic workflow to atom-map reactions and standardize inconsistencies in representation across entries. **D**) Distribution of the 599 reconstructed reactions across EC classes, showing substantial class imbalance. **E**) Mechanistic steps of the CYP aliphatic hydroxylation of P450-BM3 enzymes (M-CSA entry 699) [26]. Fluxion is fine-tuned using publicly available CYP reaction data to generate sequence-conditioned distributions over alternative selectivity-determining hydrogen atom transfers for step five.

**Figure S2:**
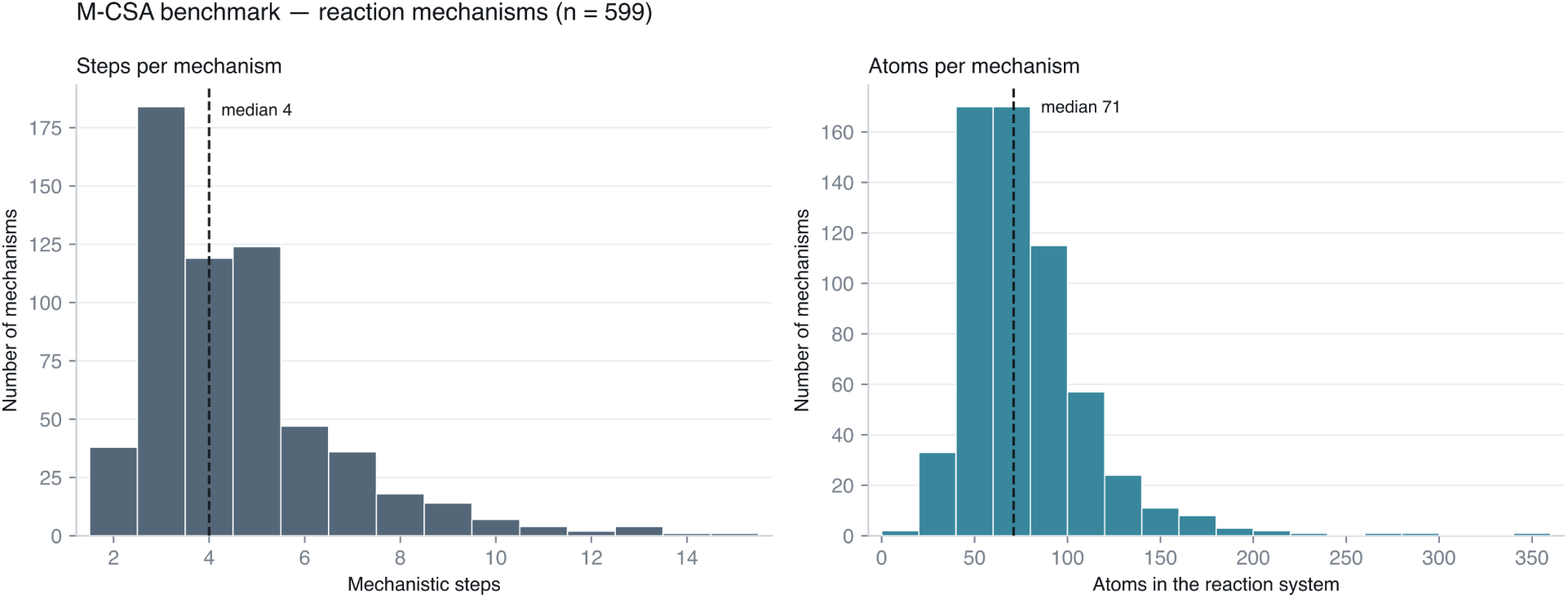
Distribution of mechanism step length and size across the reconstructed M-CSA dataset. Distribution of the number of mechanistic steps (left) and total number of atoms (right) across the 599 reconstructed M-CSA mechanisms. Both distributions are right-skewed, with a small number of substantially longer or larger mechanisms.

**Figure S3:**
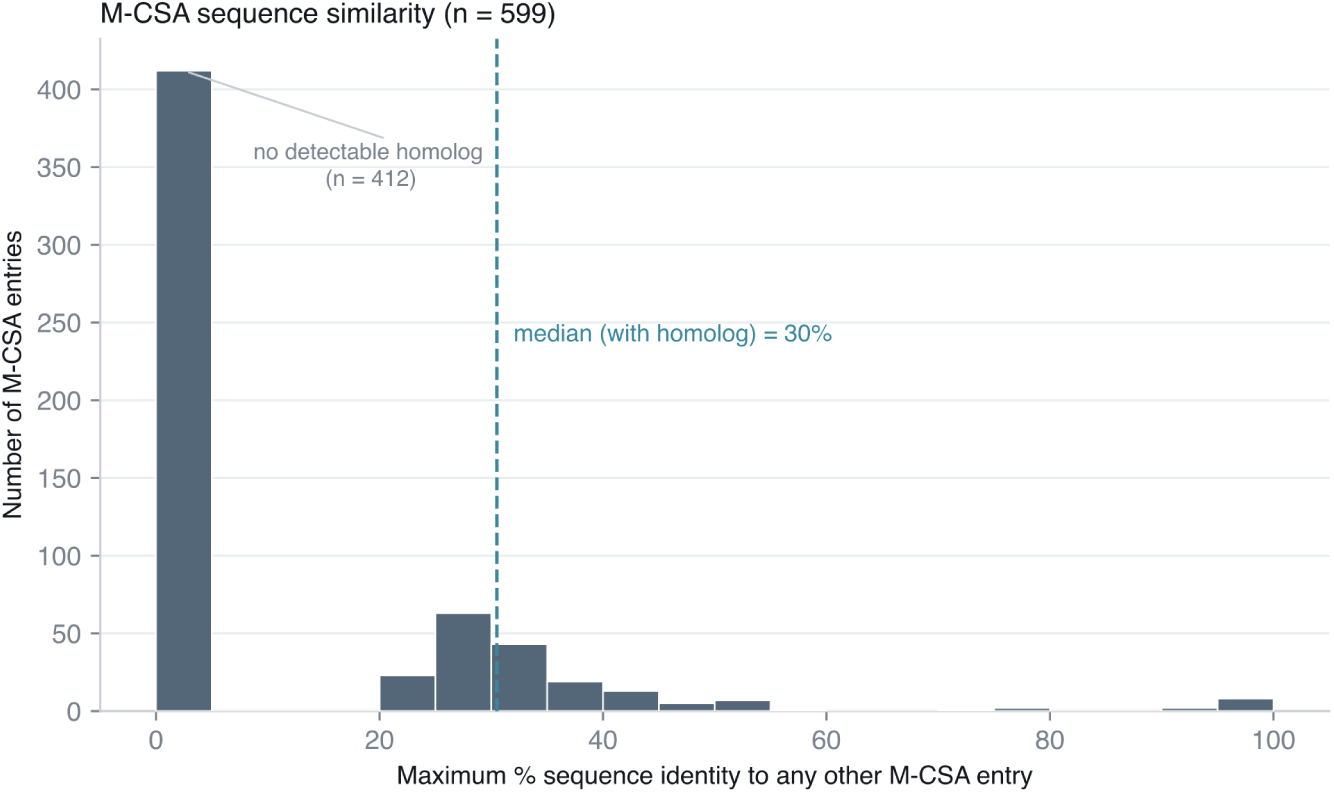
Sequence redundancy within the reconstructed M-CSA dataset. Distribution of the maximum sequence identity of each of the 599 M-CSA enzyme sequences to any other entry in the dataset. Most entries (412/599) have no detectable homologue within M-CSA; among entries with a detectable homologue, the median maximum sequence identity is 30%. Thus, most mechanisms are represented by isolated or weakly related enzyme sequences rather than multiple closely related members of the same enzyme family.

**Figure S4:**
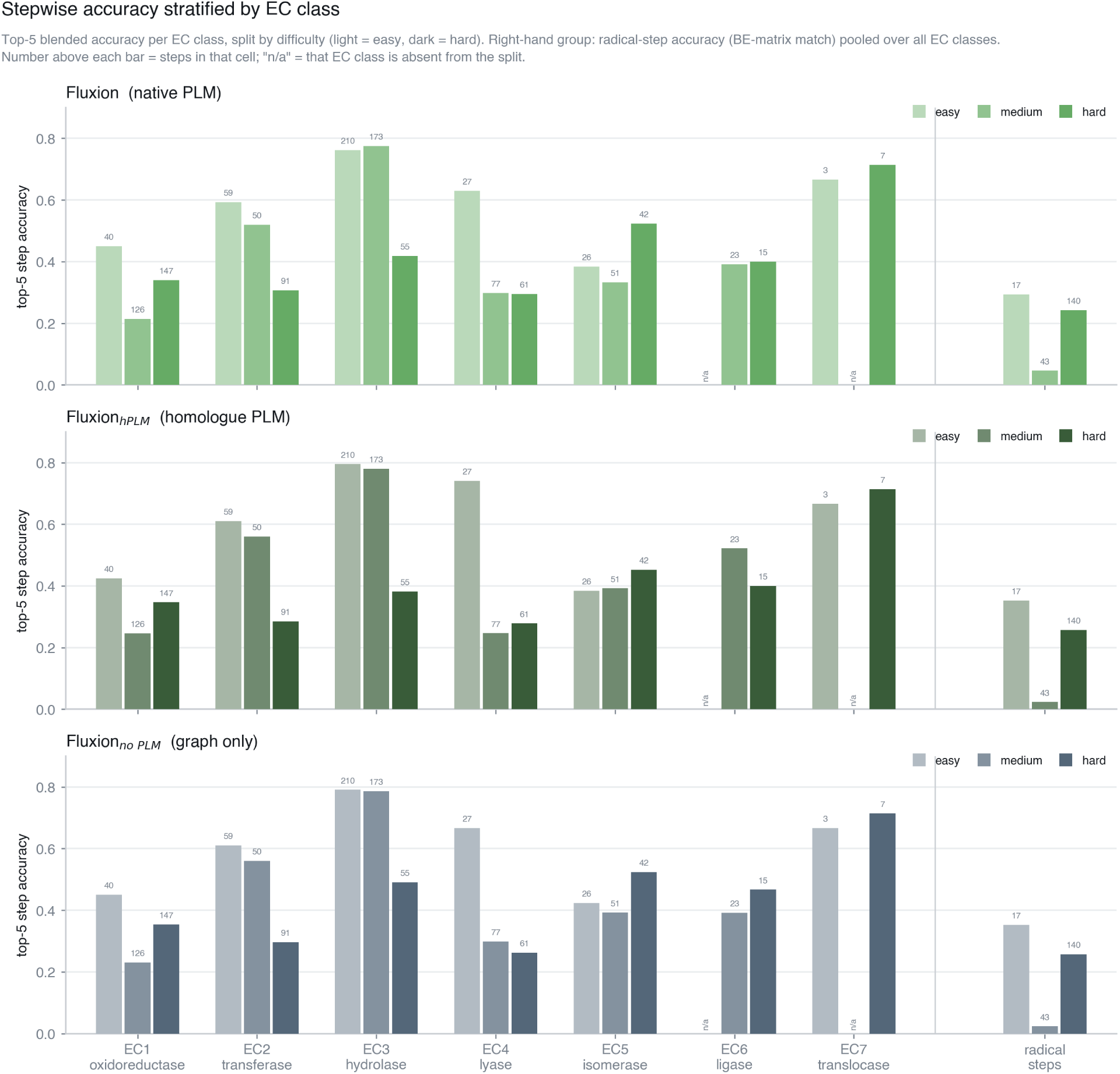
Stepwise mechanism prediction accuracy stratified by EC class. Top-5 stepwise accuracy is shown for the easy, medium, and hard M-CSA benchmark splits across EC classes for Fluxion with native PLM conditioning, homologue PLM conditioning, or no PLM conditioning. Numbers above bars indicate the number of mechanistic steps evaluated in each group (the denominator of the accuracy). Radical-step accuracy is shown separately, pooled across EC classes.

**Figure S5:**
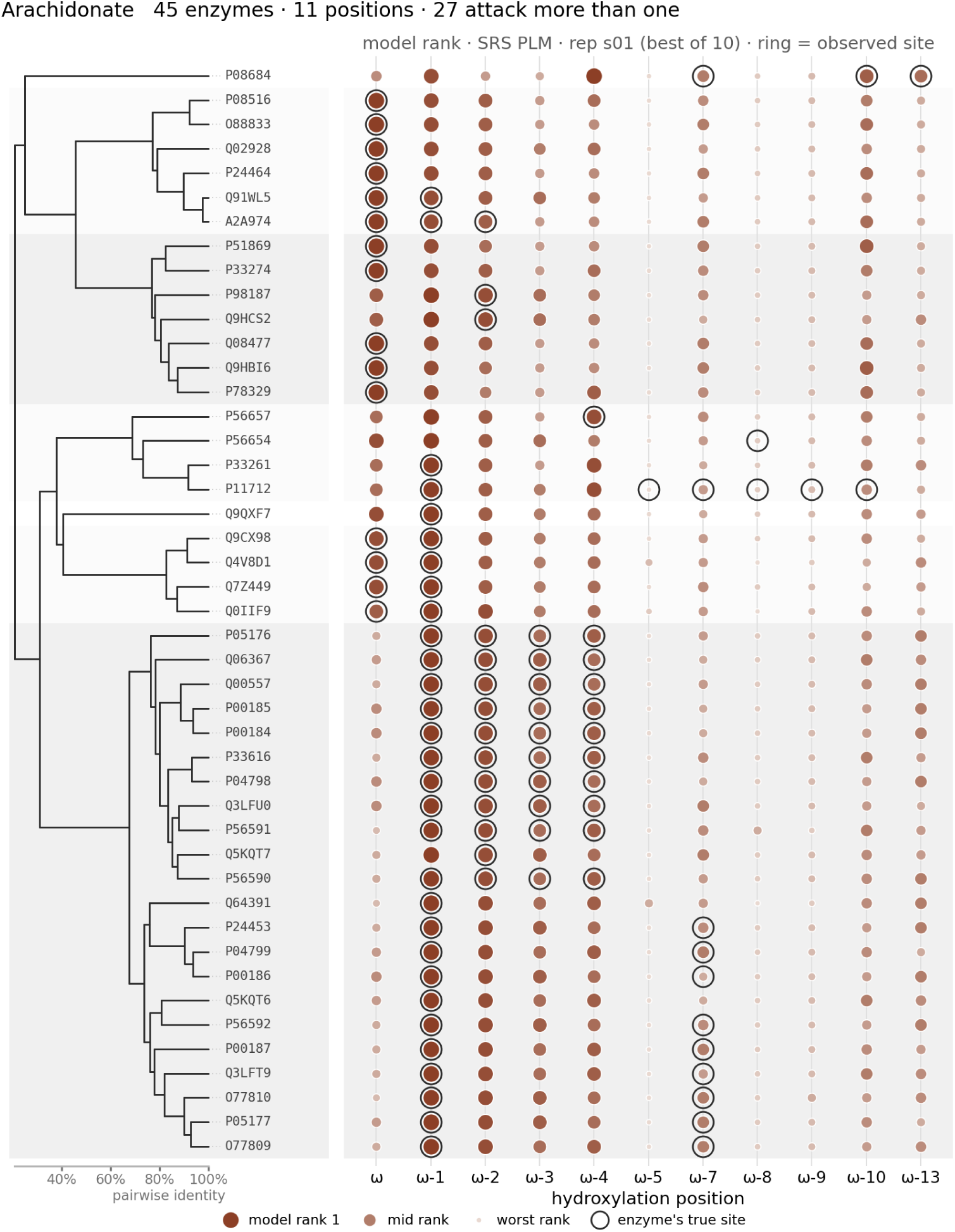
Sequence-conditioned regioselectivity predictions for arachidonate. Predicted hydroxylation-site rankings for 45 CYP enzymes acting on arachidonate. The dendrogram (left) shows pairwise sequence relationships between enzymes. For each enzyme, candidate hydroxylation positions are ranked by Fluxion using the SRS-conditioned model (right), with larger and darker points indicating higher predicted rank. Black rings denote experimentally observed hydroxylation sites. Twenty-seven enzymes were experimentally observed to hydroxylate more than one position.

**Figure S6:**
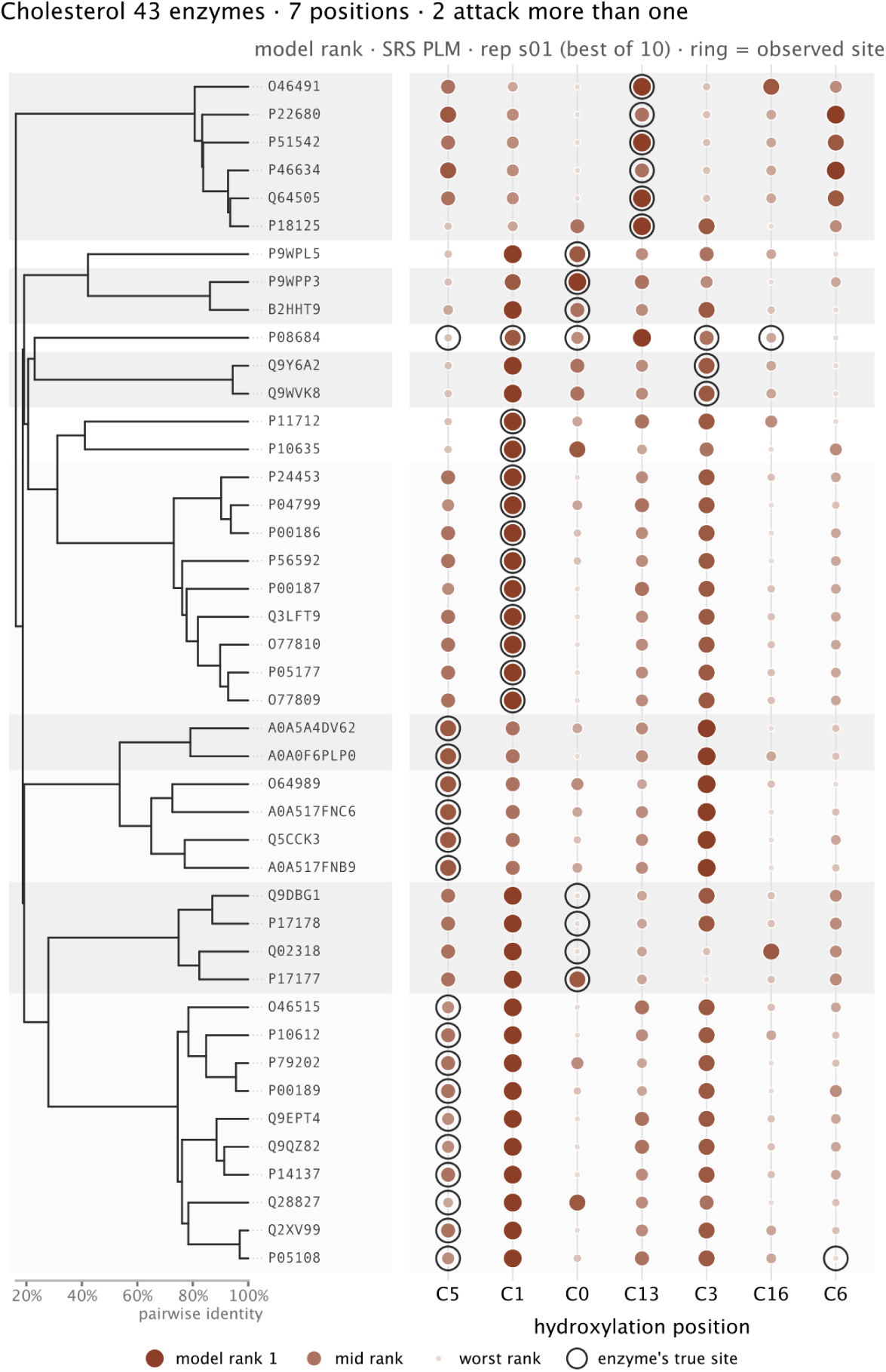
Sequence-conditioned regioselectivity predictions for cholesterol. Predicted hydroxylation-site rankings for 43 CYP enzymes acting on cholesterol. The dendrogram (left) shows pairwise sequence relationships between enzymes. For each enzyme, candidate hydroxylation positions are ranked by Fluxion using the SRS- conditioned model (right), with larger and darker points indicating higher predicted rank. Black rings denote experimentally observed hydroxylation sites. Only two enzymes were experimentally observed to hydroxylate more than one position.

**Figure S7:**
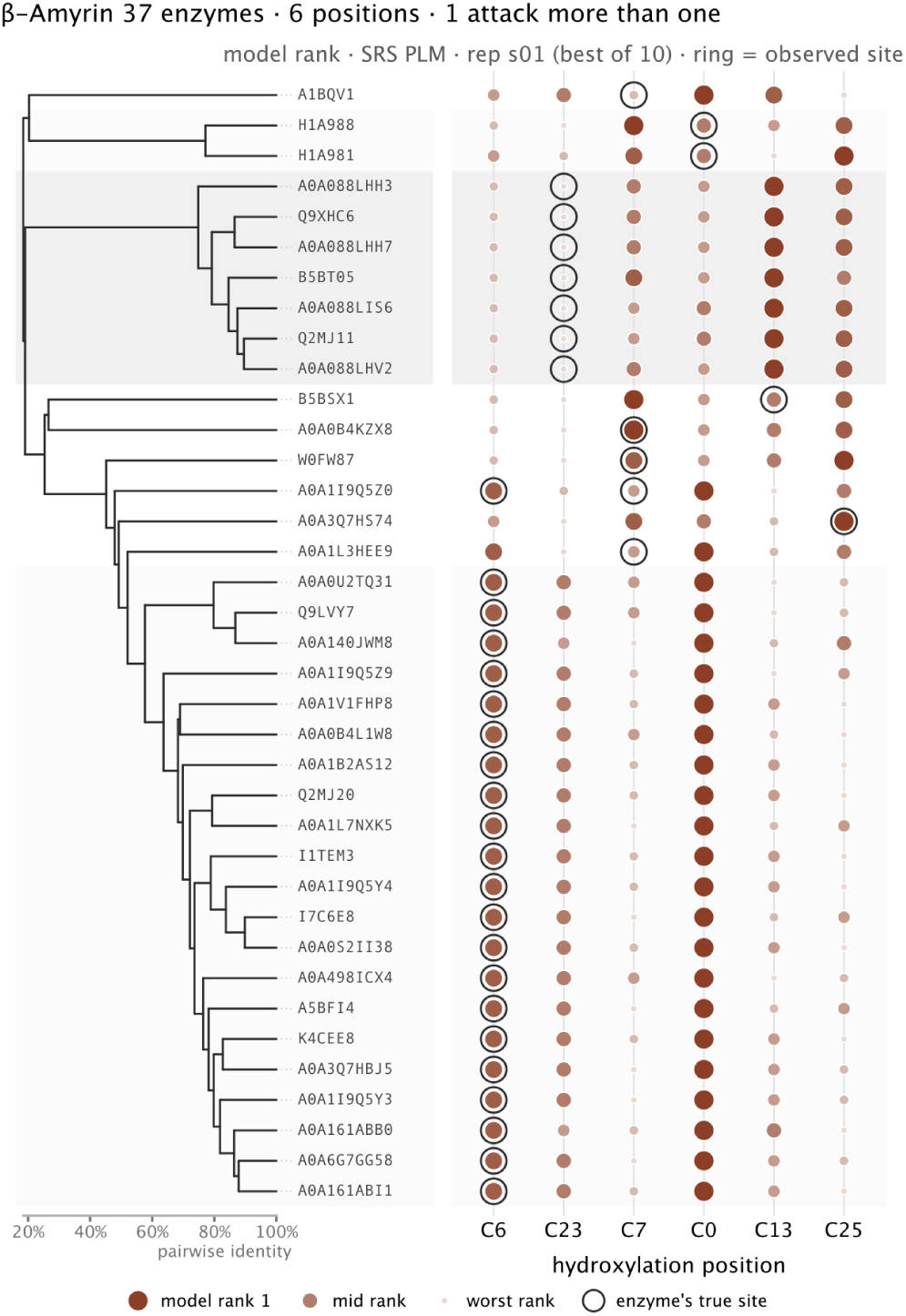
Sequence-conditioned regioselectivity predictions for β-amyrin. Predicted hydroxylation-site rankings for 37 CYP enzymes acting on *β*-amyrin. The dendrogram (left) shows pairwise sequence relationships between enzymes. For each enzyme, candidate hydroxylation positions are ranked by Fluxion using the SRS-conditioned model (right), with larger and darker points indicating higher predicted rank. Black rings denote experimentally observed hydroxylation sites. Only one enzyme was experimentally observed to hydroxylate more than one position.

**Figure S8:**
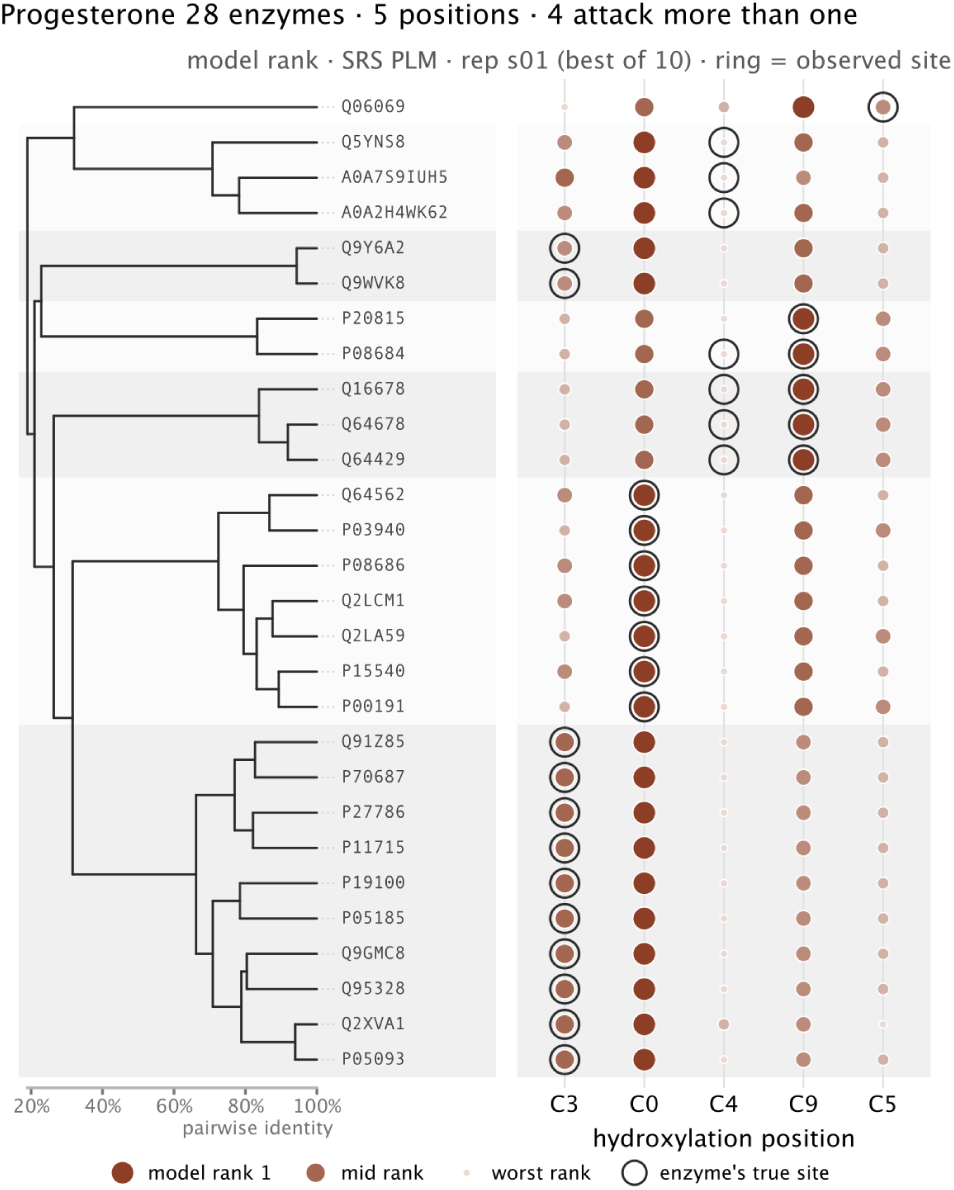
Sequence-conditioned regioselectivity predictions for progesterone. Predicted hydroxylation-site rankings for 28 CYP enzymes acting on progesterone. The dendrogram (left) shows pairwise sequence relationships between enzymes. For each enzyme, candidate hydroxylation positions are ranked by Fluxion using the SRS-conditioned model (right), with larger and darker points indicating higher predicted rank. Black rings denote experimentally observed hydroxylation sites. Four enzymes were experimentally observed to hydroxylate more than one position.

**Figure S9:**
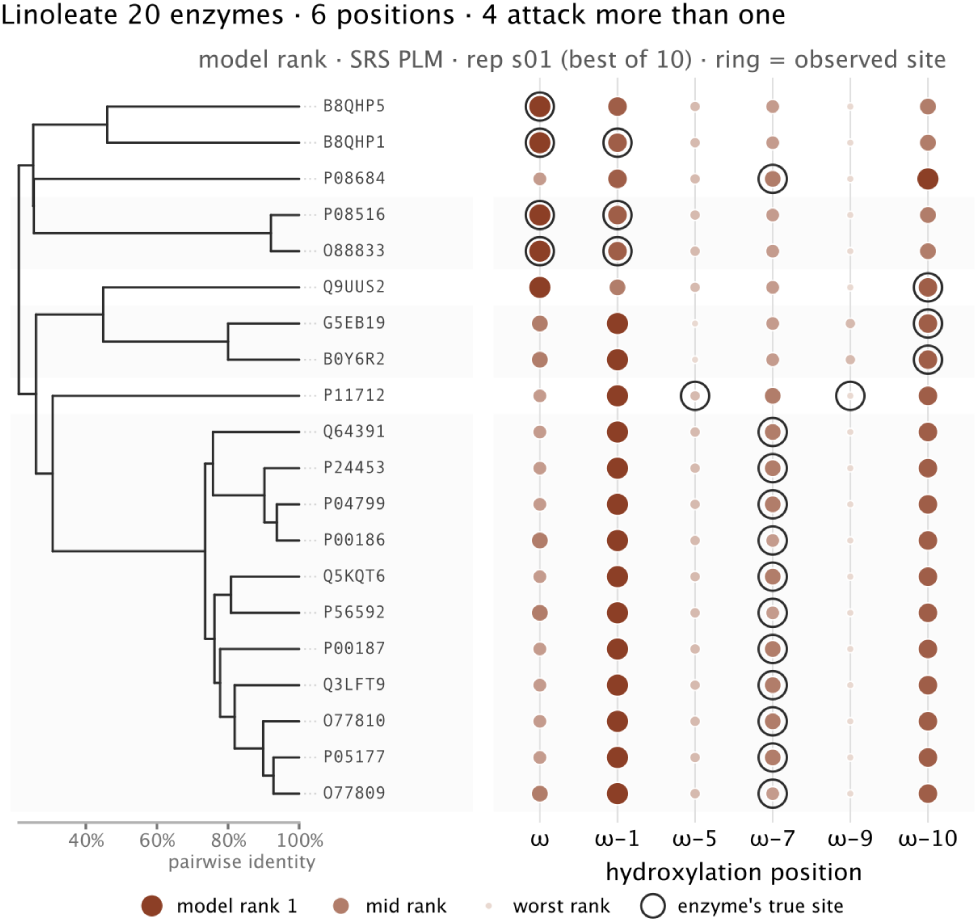
Sequence-conditioned regioselectivity predictions for linoleate. Predicted hydroxylation-site rankings for 20 CYP enzymes acting on linoleate. The dendrogram (left) shows pairwise sequence relationships between enzymes. For each enzyme, candidate hydroxylation positions are ranked by Fluxion using the SRS-conditioned model (right), with larger and darker points indicating higher predicted rank. Black rings denote experimentally observed hydroxylation sites. Four enzymes were experimentally observed to hydroxylate more than one position.

**Figure S10:**
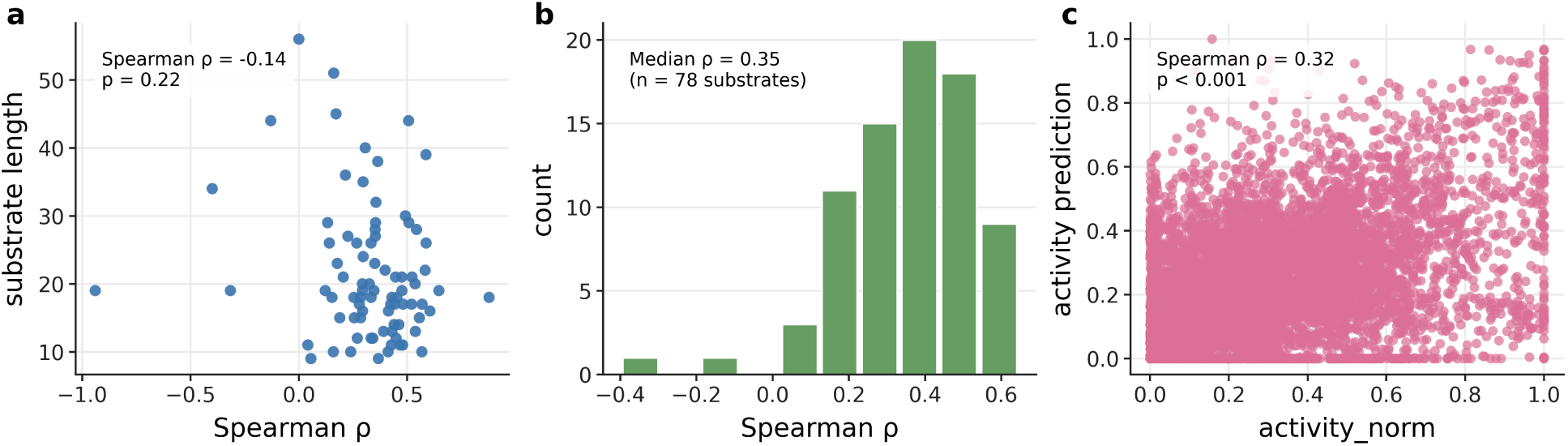
**A**) Correlation between the size of the substrate and the within substrate correlation for the esterase dataset from Fluxion model trained on both USPTO and EnzymeMap followed by finetuning. **B**) Within substrate Spearman’s *ρ* between predicted and true activity for active enzymes from the esterase dataset using the Fluxion model trained on both USPTO and EnzymeMap followed by finetuning **C**) Correlation across the esterase dataset between predicted and true normalized activity (log_2_ followed by standard scaling and min-max) using the Fluxion model trained on both EnzymeMap and USPTO followed by finetuning.

**Figure S11:**
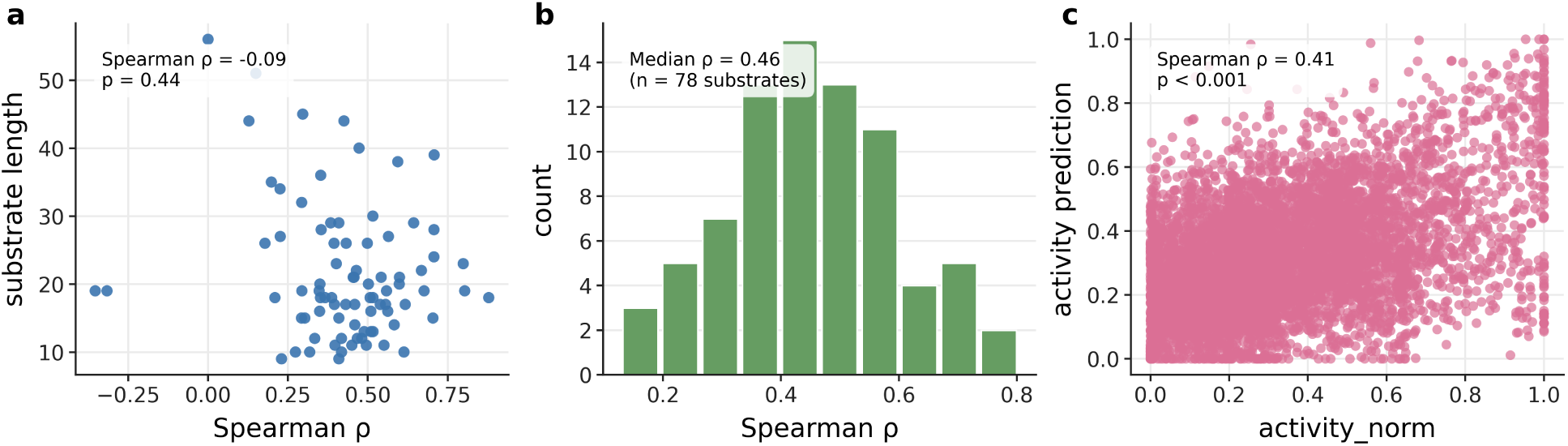
**A**) Correlation between the size of the substrate and the within substrate correlation for the esterase dataset from Fluxion model trained on only USPTO followed by finetuning. **B**) Within substrate Spearman’s *ρ* between predicted and true activity for active enzymes from the esterase dataset using the Fluxion model trained on only USPTO followed by finetuning. **C**) Correlation across the esterase dataset between predicted and true normalized activity (log_2_ followed by standard scaling and min-max) using the Fluxion model trained on only USPTO followed by finetuning.

**Figure S12:**
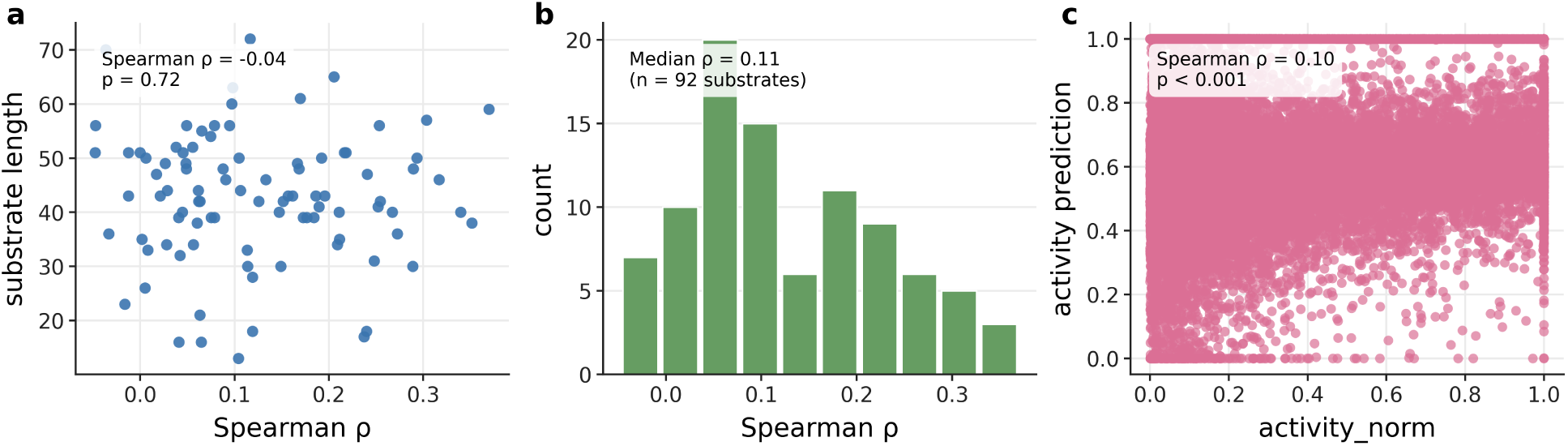
**A**) Correlation between the size of the substrate and the within substrate correlation for the phosphatase dataset. **B**) Within substrate Spearman’s *ρ* between predicted and true activity for active enzymes from the phosphatase dataset. **C**) Correlation across the phosphatase dataset between predicted and true normalized activity (log_2_ followed by standard scaling and min-max).

**Figure S13:**
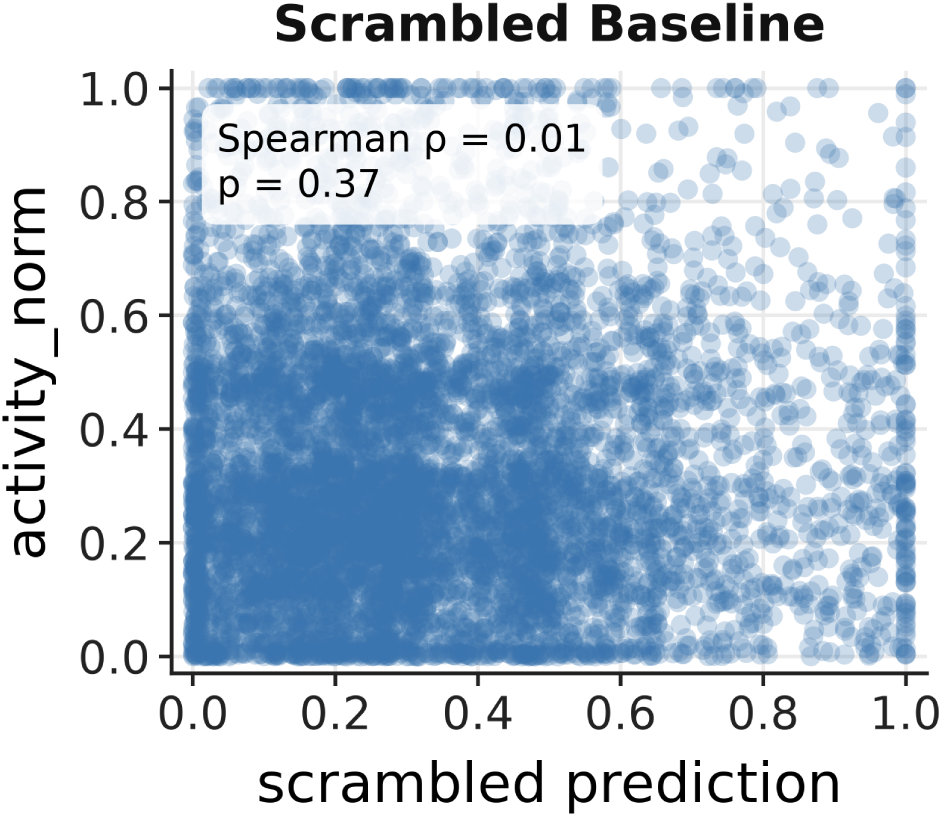
**A**) Scrambled baseline shows that random mixing of the esterase dataset would not show a correlation between the predicted and true activity.

**Figure S14:**
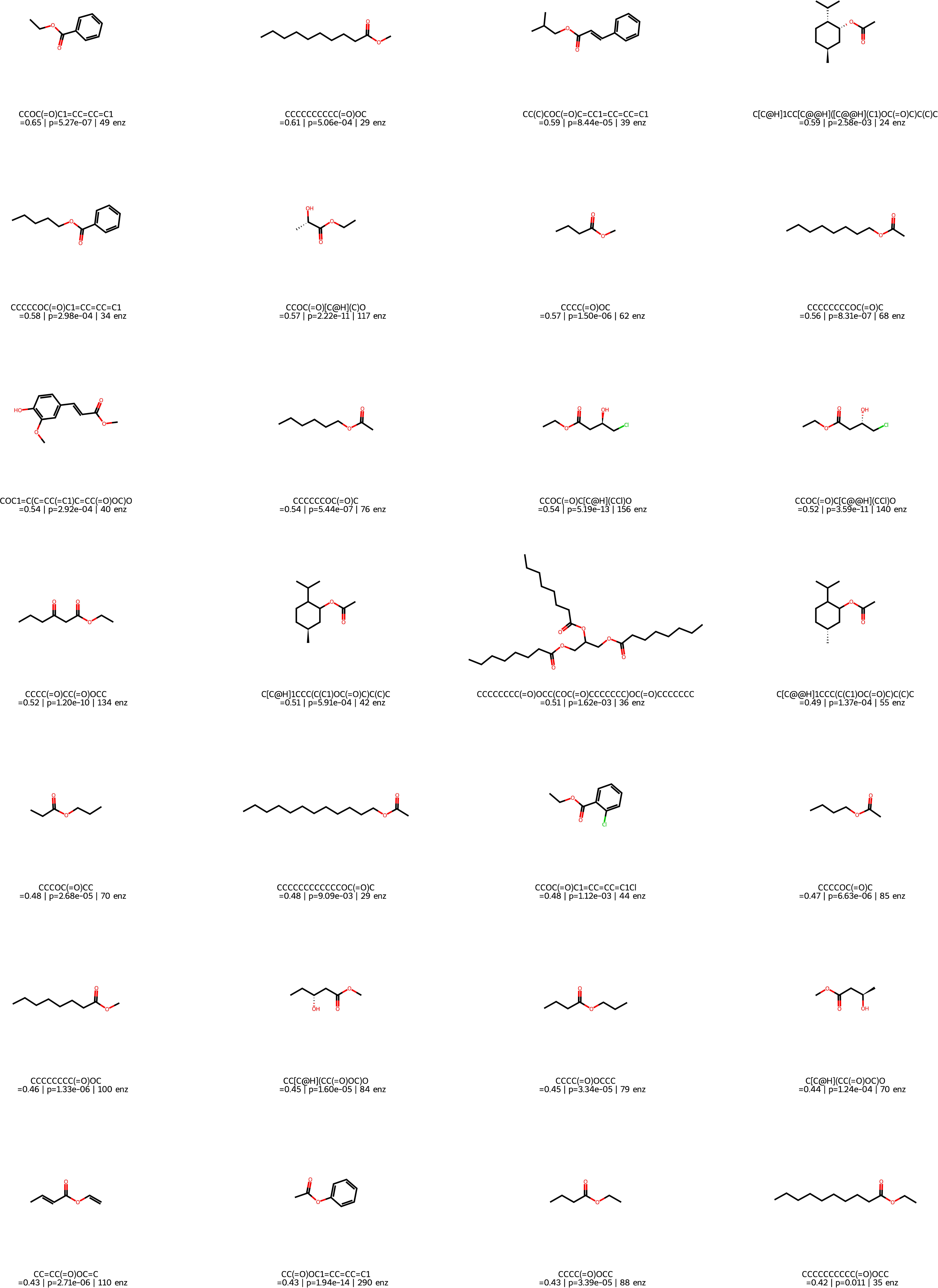
Held out substrates from the esterase dataset (all splits), the number of non zero enzymes recorded for this substrate is reported along side the Spearman’s ρ and uncorrected p-value.

**Figure S15:**
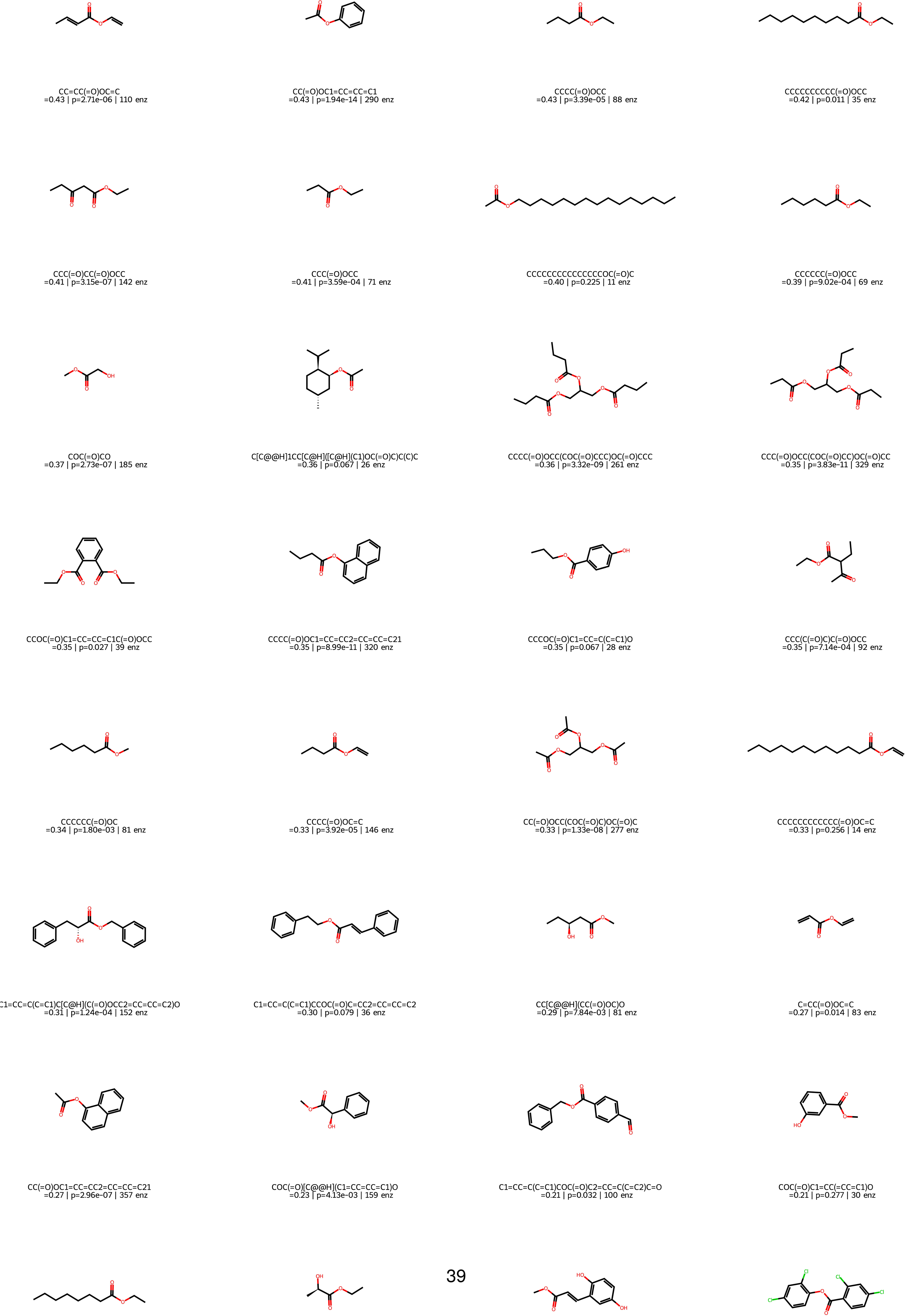
Held out substrates from the esterase dataset (all splits), the number of non zero enzymes recorded for this substrate is reported along side the Spearman’s ρ and uncorrected p-value.

**Figure S16:**
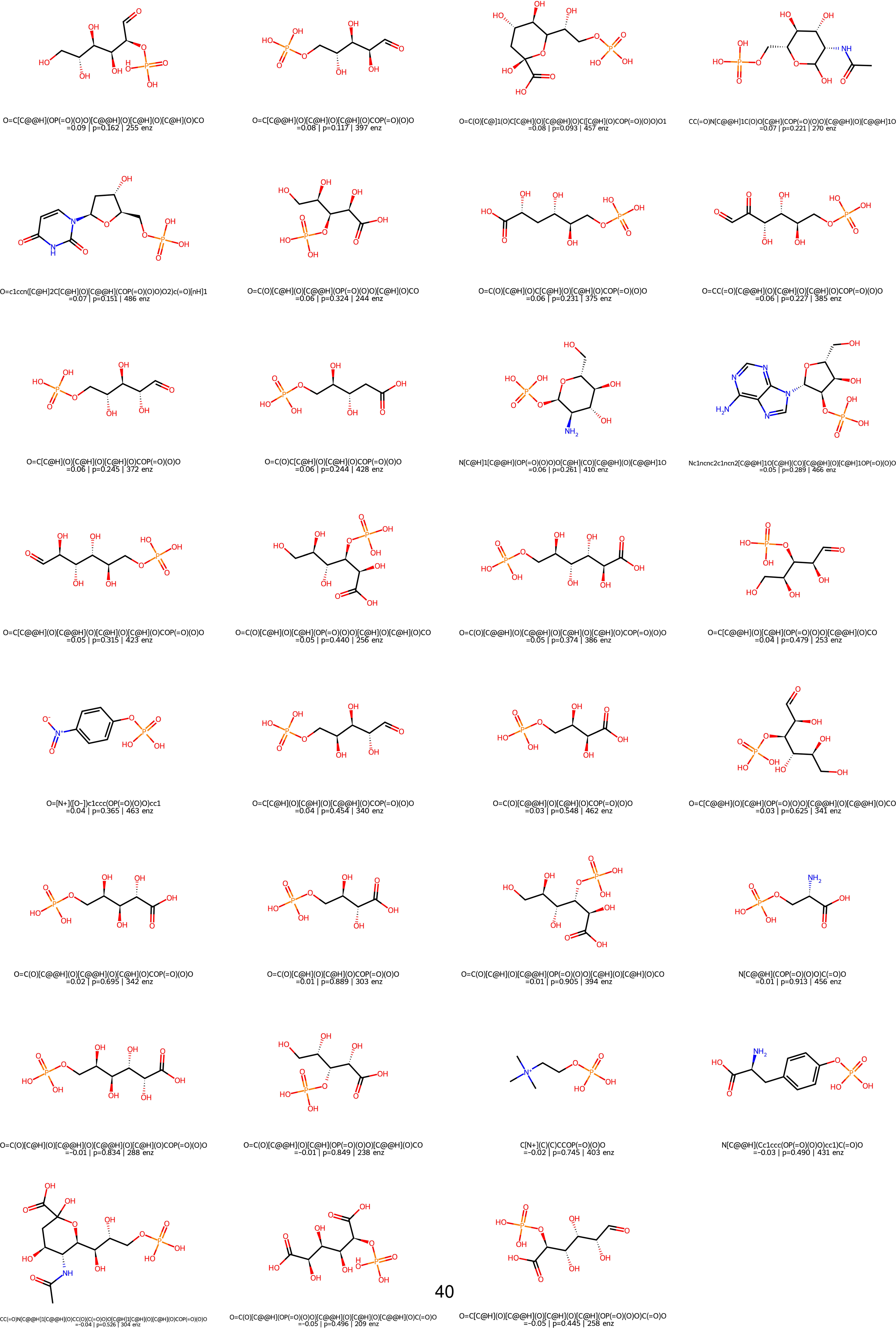
Held out substrates from the phosphatase dataset (all splits), the number of non zero enzymes recorded for this substrate is reported along side the Spearman’s ρ and uncorrected p-value.

**Figure S17:**
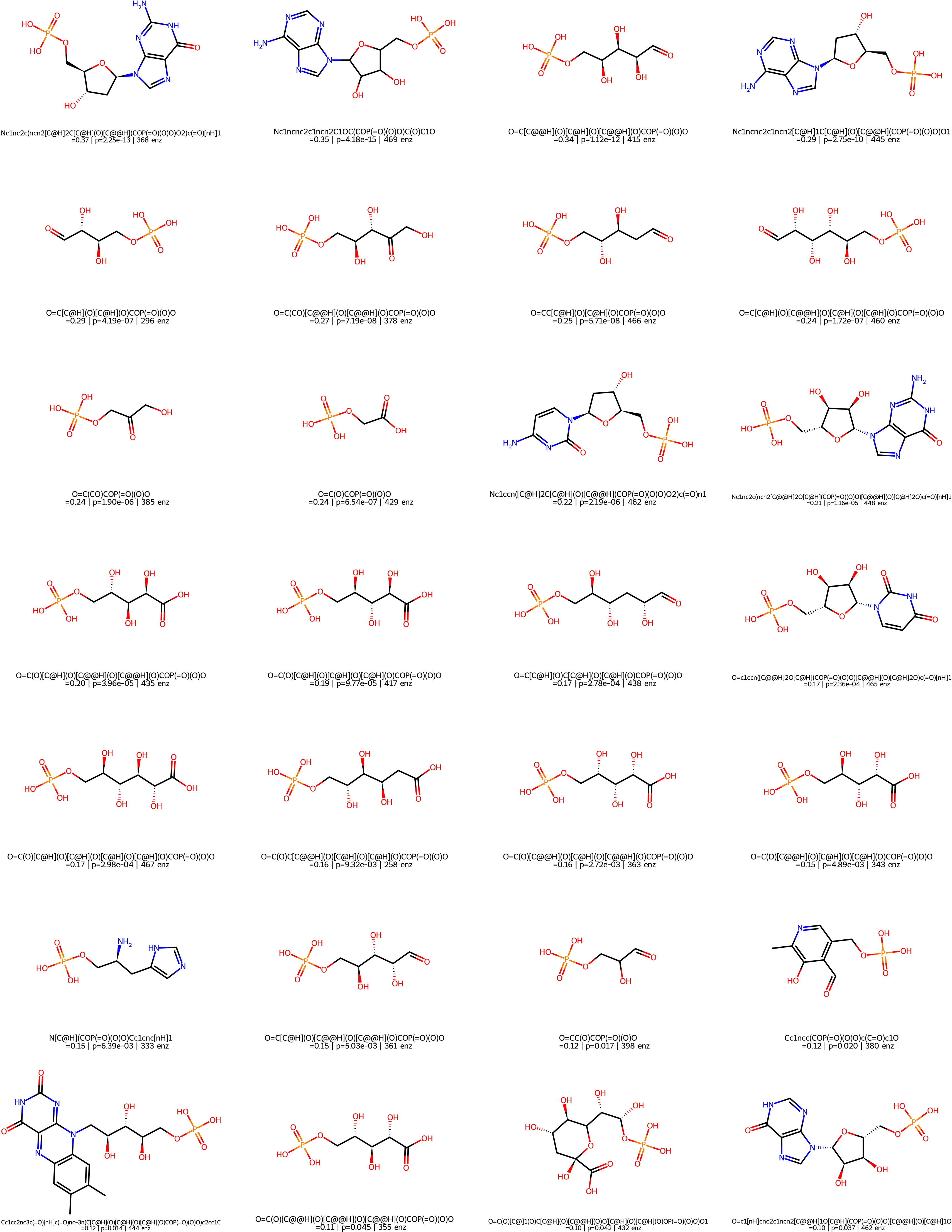
Held out substrates from the phosphatase dataset (all splits), the number of non zero enzymes recorded for this substrate is reported along side the Spearman’s ρ and uncorrected p-value.

**Figure S18:**
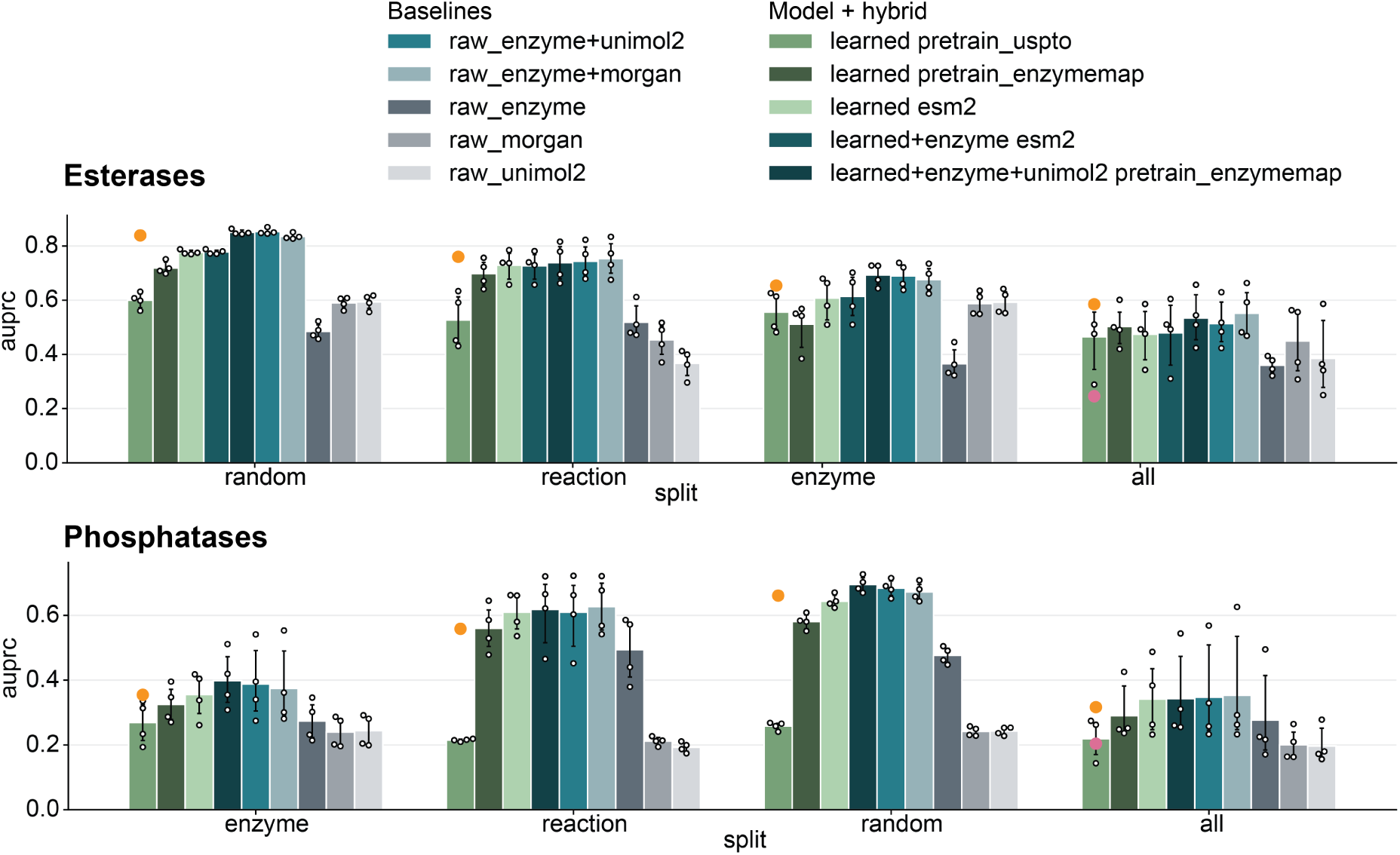
Ablations to see what information is captured by the model as compared with different baselines.

**Figure S19:**
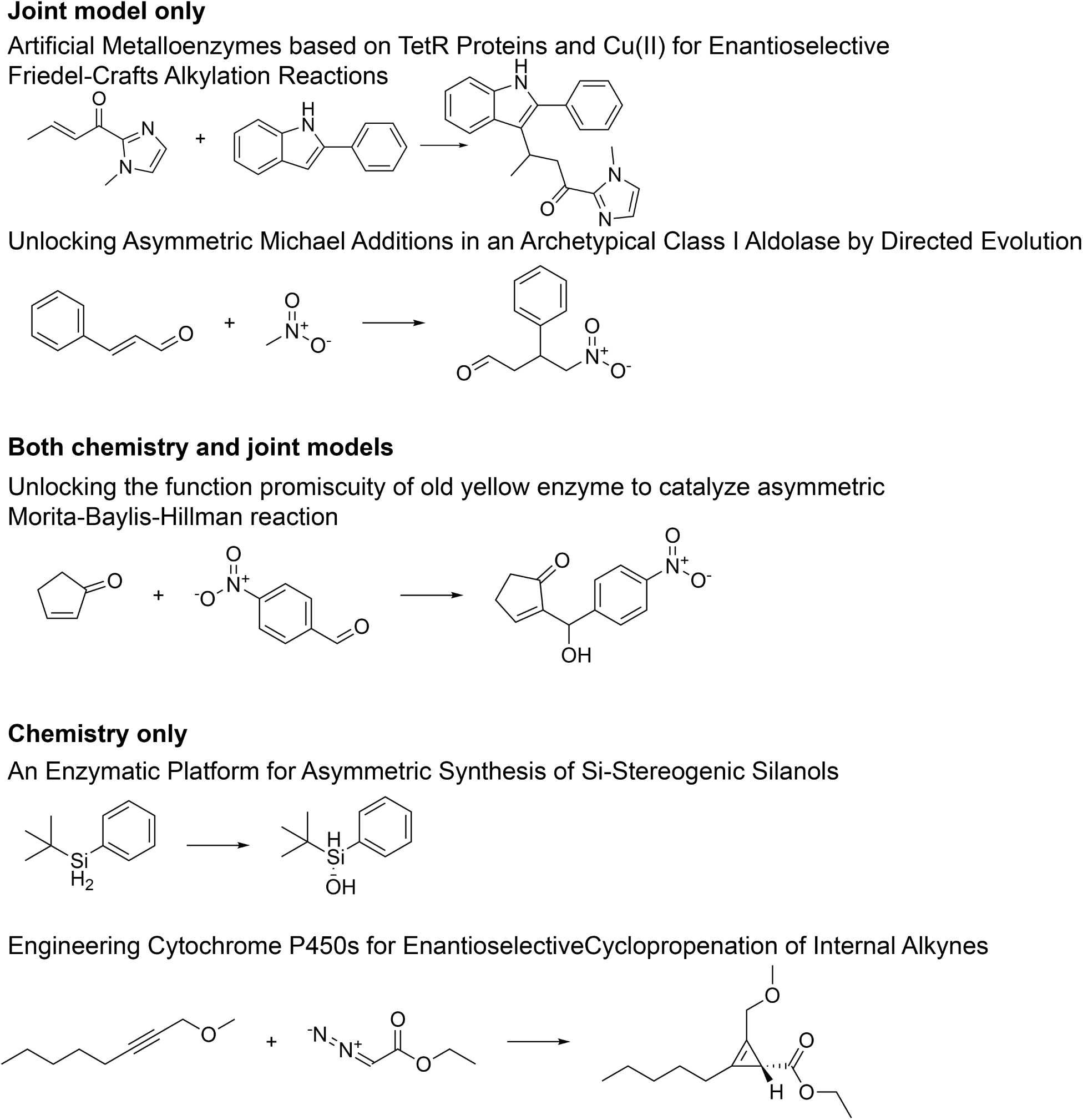
Select reactions that are either shared, or generated only by one of the trained Fluxion models.

## References

[1] Frances H. Arnold. Directed evolution: Bringing new chemistry to life. Angewandte Chemie International Edition, 57(16): 4143–4148, 2018. ISSN 1521-3773. doi: 10.1002/anie.201708408.

[2] Julia C. Reisenbauer, Kathleen M. Sicinski, and Frances H. Arnold. Catalyzing the future: recent advances in chemical synthesis using enzymes. Current Opinion in Chemical Biology, 83:102536, December 2024. ISSN 1367-5931. doi: 10.1016/j.cbpa.2024.102536.

[3] Alexandra E. Paton, Daniil A. Boiko, Jonathan C. Perkins, Nicholas I. Cemalovic, Thiago Reschützegger, Gabe Gomes, and Alison R. H. Narayan. Connecting chemical and protein sequence space to predict biocatalytic reactions. Nature, 646(8083): 108–116, October 2025. ISSN 1476-4687. doi: 10.1038/s41586-025-09519-5.

[4] Haiyang Cui, Yufeng Su, Tanner J. Dean, Tianhao Yu, Zhengyi Zhang, Jian Peng, Diwakar Shukla, and Huimin Zhao. Enzyme specificity prediction using cross-attention graph neural networks. Nature, 647(8090):639–647, November 2025. ISSN 1476-4687. doi: 10.1038/s41586-025-09697-2.

[5] Peter G. Mikhael, Itamar Chinn, and Regina Barzilay. Clipzyme: Reaction-conditioned virtual screening of enzymes. (arXiv:2402.06748), February 2024. doi: 10.48550/arXiv.2402.06748. URL http://arxiv.org/abs/2402.06748. arXiv:2402.06748 [q-bio].

[6] Jason W. Rocks, Dat P. Truong, Dmitrij Rappoport, Samuel Maddrell-Mander, Daniel A. Martin-Alarcon, Toni M. Lee, Steven Crossan, and Joshua E. Goldford. Dual-encoder contrastive learning accelerates enzyme discovery. Proceedings of the National Academy of Sciences, 123(12):e2520070123, March 2026. doi: 10.1073/pnas.2520070123.

[7] Yong Liu, Chenqing Hua, Menglong Xu, Tao Zeng, Jiahua Rao, Zhongyue Zhang, Ruibo Wu, Jing-Ke Weng, Connor W. Coley, and Shuangjia Zheng. A geometric foundation model for enzyme retrieval with evolutionary insights. Nature Catalysis, 9(2):148–160, February 2026. ISSN 2520-1158. doi: 10.1038/s41929-026-01478-y.

[8] Alexander Kroll, Sahasra Ranjan, Martin K. M. Engqvist, and Martin J. Lercher. A general model to predict small molecule substrates of enzymes based on machine and deep learning. Nature Communications, 14(1):2787, May 2023. ISSN 2041-1723. doi: 10.1038/s41467-023-38347-2.

[9] Elizabeth H. Mahood, Natália Komorníková, Tomáš Pluskal, and Pranam Chatterjee. Rethinking benchmarks and models for enzyme specificity prediction. (arXiv:2607.05084), July 2026. doi: 10.48550/arXiv.2607.05084. URL http://arxiv.org/abs/2607.05084. arXiv:2607.05084 [q-bio.BM].

[10] Jason Yang, Ariane Mora, Shengchao Liu, Bruce James Wittmann, Anima Anandkumar, Frances H. Arnold, and Yisong Yue. Care: a benchmark suite for the classification and retrieval of enzymes. November 2024. URL https://openreview.net/forum?id=PFwlw9bnAr#discussion.

[11] Joseph L. Watson, David Juergens, Nathaniel R. Bennett, Brian L. Trippe, Jason Yim, Helen E. Eisenach, Woody Ahern, Andrew J. Borst, Robert J. Ragotte, Lukas F. Milles, Basile I. M. Wicky, Nikita Hanikel, Samuel J. Pellock, Alexis Courbet, William Sheffler, Jue Wang, Preetham Venkatesh, Isaac Sappington, Susana Vázquez Torres, Anna Lauko, Valentin De Bortoli, Emile Mathieu, Sergey Ovchinnikov, Regina Barzilay, Tommi S. Jaakkola, Frank DiMaio, Minkyung Baek, and David Baker. De novo design of protein structure and function with rfdiffusion. Nature, 620(7976):1089–1100, August 2023. ISSN 1476-4687. doi: 10.1038/s41586-023-06415-8.

[12] Anna Lauko, Samuel J. Pellock, Kiera H. Sumida, Ivan Anishchenko, David Juergens, Woody Ahern, Jihun Jeung, Alex Shida, Andrew Hunt, Indrek Kalvet, Christoffer Norn, Ian R. Humphreys, Cooper Jamieson, Rohith Krishna, Yakov Kipnis, Alex Kang, Evans Brackenbrough, Asim K. Bera, Banumathi Sankaran, K. N. Houk, and David Baker. Computational design of serine hydrolases. Science, 0(0):eadu2454, February 2025. doi: 10.1126/science.adu2454.

[13] Jarrid Rector-Brooks, Théophile Lambert, Marta Skreta, Daniel Roth, Yueming Long, Zi-Qi Li, Xi Zhang, Miruna Cretu, Francesca-Zhoufan Li, Tanvi Ganapathy, Emily Jin, Avishek Joey Bose, Jason Yang, Kirill Neklyudov, Yoshua Bengio, Alexander Tong, Frances H. Arnold, and Cheng-Hao Liu. General multimodal protein design enables dna-encoding of chemistry. (arXiv:2604.05181), April 2026. doi: 10.48550/arXiv.2604.05181. URL http://arxiv.org/abs/2604.05181. arXiv:2604.05181 [cs.LG].

[14] Emily Jin, Andrei Cristian Nica, Mikhail Galkin, Jarrid Rector-Brooks, Kin Long Kelvin Lee, Santiago Miret, Frances H. Arnold, Michael Bronstein, Avishek Joey Bose, Alexander Tong, and Cheng-Hao Liu. Oxtal: An all-atom diffusion model for organic crystal structure prediction. (arXiv:2512.06987), April 2026. doi: 10.48550/arXiv.2512.06987. URL http://arxiv.org/abs/2512.06987. arXiv:2512.06987 [cs.LG].

[15] Joonyoung F. Joung, Mun Hong Fong, Nicholas Casetti, Jordan P. Liles, Ne S. Dassanayake, and Connor W. Coley. Electron flow matching for generative reaction mechanism prediction. Nature, 645(8079):115–123, September 2025. ISSN 1476-4687. doi: 10.1038/s41586-025-09426-9.

[16] Daniel Probst, Matteo Manica, Yves Gaetan Nana Teukam, Alessandro Castrogiovanni, Federico Paratore, and Teodoro Laino. Biocatalysed synthesis planning using data-driven learning. Nature communications, 13(1):964, 2022.

[17] Lun-Hsin Kuo, Jason Yang, and Frances Arnold. Ezsolver: Template-free prediction of polar enzymatic mechanisms via bidirectional flow matching and search. bioRxiv, pages 2026–07, 2026.

[18] Guan-Horng Liu, Arash Vahdat, De-An Huang, Evangelos A. Theodorou, Weili Nie, and Anima Anandkumar. I^2^sb: Image-to-image schrödinger bridge, 2023. URL https://arxiv.org/abs/2302.05872.

[19] Antonio J. M. Ribeiro, Ioannis G. Riziotis, Neera Borkakoti, Pedro A. Fernandes, Maria J. Ramos, and Janet M. Thornton. Measuring catalytic mechanism similarity – a new approach to study enzyme function and evolution. The FEBS Journal, 292 (16):4200–4210, 2025. doi: 10.1111/febs.70106. URL https://febs.onlinelibrary.wiley.com/doi/abs/10.1111/febs.70106.

[20] Austin D Hartley, Vikas Upadhyay, Veda Sheersh Boorla, and Costas D Maranas. Mechfind: a computational framework for de novo prediction of enzyme mechanisms. Nature communications, 17(1):3903, 2026.

[21] James Dugundji and Ivar Ugi. An algebraic model of constitutional chemistry as a basis for chemical computer programs. In Computers in Chemistry, page 19–64, Berlin, Heidelberg, 1973. Springer. ISBN 978-3-540-38510-3. doi: 10.1007/BFb0051317.

[22] Zeming Lin, Halil Akin, Roshan Rao, Brian Hie, Zhongkai Zhu, Wenting Lu, Nikita Smetanin, Robert Verkuil, Ori Kabeli, Yaniv Shmueli, Allan dos Santos Costa, Maryam Fazel-Zarandi, Tom Sercu, Salvatore Candido, and Alexander Rives. Evolutionary-scale prediction of atomic-level protein structure with a language model. Science, 379(6637):1123–1130, March 2023. doi: 10.1126/science.ade2574.

[23] Yili Shen and Xiangliang Zhang. Driving reaction trajectories via latent flow matching, 2026. URL https://arxiv.org/abs/2602.10476.

[24] Haitao Lin, Junjie Wang, Zhifeng Gao, Xiaohong Ji, Rong Zhu, Linfeng Zhang, Guolin Ke, and Weinan E. Synbridge: Bridging reaction states via discrete flow for bidirectional reaction prediction, 2025. URL https://arxiv.org/abs/2507.08475.

[25] Robin Yadav, Qi Yan, Guy Wolf, Joey Bose, and Renjie Liao. Retro synflow: Discrete flow-matching for accurate and diverse single-step retrosynthesis. Advances in Neural Information Processing Systems, 38:55229–55258, 2026.

[26] António J M Ribeiro, Gemma L Holliday, Nicholas Furnham, Jonathan D Tyzack, Katherine Ferris, and Janet M Thornton. Mechanism and catalytic site atlas (m-csa): a database of enzyme reaction mechanisms and active sites. Nucleic Acids Research, 46(D1):D618–D623, January 2018. ISSN 0305-1048, 1362-4962. doi: 10.1093/nar/gkx1012.

[27] Jonathan D Tyzack and Johannes Kirchmair. Computational methods and tools to predict cytochrome p450 metabolism for drug discovery. Chemical biology & drug design, 93(4):377–386, 2019.

[28] Vladimir Porokhin, Li-Ping Liu, and Soha Hassoun. Using graph neural networks for site-of-metabolism prediction and its applications to ranking promiscuous enzymatic products. Bioinformatics, 39(3):btad089, 2023.

[29] Xuhai Huang, Jiamin Chang, and Boxue Tian. Glmcyp: a deep learning-based method for cyp450-mediated reaction site prediction. Journal of Chemical Information and Modeling, 65(5):2322–2335, 2025.

[30] Jiamin Chang, Xuhai Huang, and Boxue Tian. Metacyp: a unified framework for prediction of cytochrome p450 metabolic sites and reaction types via multimodal deep learning. Frontiers in Chemistry, 14:1869559, 2026.

[31] John Jumper, Richard Evans, Alexander Pritzel, Tim Green, Michael Figurnov, Olaf Ronneberger, Kathryn Tunyasuvunakool, Russ Bates, Augustin Žídek, Anna Potapenko, Alex Bridgland, Clemens Meyer, Simon A. A. Kohl, Andrew J. Ballard, Andrew Cowie, Bernardino Romera-Paredes, Stanislav Nikolov, Rishub Jain, Jonas Adler, Trevor Back, Stig Petersen, David Reiman, Ellen Clancy, Michal Zielinski, Martin Steinegger, Michalina Pacholska, Tamas Berghammer, Sebastian Bodenstein, David Silver, Oriol Vinyals, Andrew W. Senior, Koray Kavukcuoglu, Pushmeet Kohli, and Demis Hassabis. Highly accurate protein structure prediction with AlphaFold. Nature, 596(7873):583–589, August 2021. ISSN 1476-4687. doi: 10.1038/s41586-021-03819-2. URL 10.1038/s41586-021-03819-2.

[32] Mónica Martínez-Martínez, Cristina Coscolín, Gerard Santiago, Jennifer Chow, Peter J. Stogios, Rafael Bargiela, Christoph Gertler, José Navarro-Fernández, Alexander Bollinger, Stephan Thies, Celia Méndez-García, Ana Popovic, Greg Brown, Tatyana N. Chernikova, Antonio García-Moyano, Gro E. K. Bjerga, Pablo Pérez-García, Tran Hai, Mercedes V. Del Pozo, Runar Stokke, Ida H. Steen, Hong Cui, Xiaohui Xu, Boguslaw P. Nocek, María Alcaide, Marco Distaso, Victoria Mesa, Ana I. Peláez, Jesús Sánchez, Patrick C. F. Buchholz, Jürgen Pleiss, Antonio Fernández-Guerra, Frank O. Glöckner, Olga V. Golyshina, Michail M. Yakimov, Alexei Savchenko, Karl-Erich Jaeger, Alexander F. Yakunin, Wolfgang R. Streit, Peter N. Golyshin, Víctor Guallar, Manuel Ferrer, and The INMARE Consortium. Determinants and prediction of esterase substrate promiscuity patterns. ACS Chemical Biology, 13(1):225–234, January 2018. ISSN 1554-8929. doi: 10.1021/acschembio.7b00996.

[33] Hua Huang, Chetanya Pandya, Chunliang Liu, Nawar F. Al-Obaidi, Min Wang, Li Zheng, Sarah Toews Keating, Miyuki Aono, James D. Love, Brandon Evans, Ronald D. Seidel, Brandan S. Hillerich, Scott J. Garforth, Steven C. Almo, Patrick S. Mariano, Debra Dunaway-Mariano, Karen N. Allen, and Jeremiah D. Farelli. Panoramic view of a superfamily of phosphatases through substrate profiling. Proceedings of the National Academy of Sciences, 112(16):E1974–E1983, April 2015. doi: 10.1073/pnas.1423570112.

[34] Samuel Goldman, Ria Das, Kevin K. Yang, and Connor W. Coley. Machine learning modeling of family wide enzyme-substrate specificity screens. PLOS Computational Biology, 18(2):e1009853, February 2022. ISSN 1553-7358. doi: 10.1371/journal.pcbi.1009853.

[35] Kai Chen and Frances H. Arnold. Engineering cytochrome p450s for enantioselective cyclopropenation of internal alkynes. Journal of the American Chemical Society, 142(15):6891–6895, March 2020. ISSN 0002-7863. doi: 10.1021/jacs.0c01313.

[36] Lei Wang, Yaoyun Wu, Jun Hu, Dejing Yin, Wanqing Wei, Jian Wen, Xiulai Chen, Cong Gao, Yiwen Zhou, Jia Liu, Guipeng Hu, Xiaomin Li, Jing Wu, Zhi Zhou, Liming Liu, and Wei Song. Unlocking the function promiscuity of old yellow enzyme to catalyze asymmetric morita-baylis-hillman reaction. Nature Communications, 15(1):5737, 2024. ISSN 2041-1723. doi: 10.1038/s41467-024-50141-2.

[37] Shunsuke Kato, Shuto Fujisawa, Yuto Adachi, Mitsuhiro Bandai, Yutaro Mori, Seiji Mori, Tomokazu Shirai, and Takashi Hayashi. Nhc-mediated radical acylation catalyzed by thiamine- and flavin-dependent enzymes. Journal of the American Chemical Society, 147(17):14837–14844, April 2025. ISSN 0002-7863. doi: 10.1021/jacs.5c04484.

[38] Chai Discovery. Zero-shot antibody design in a 24-well plate. bioRxiv, 2025. doi: 10.1101/2025.07.05.663018. URL https://www.biorxiv.org/content/early/2025/07/06/2025.07.05.663018.

[39] Yueming Long, Fatemeh Abbasinejad, Francesca-Zhoufan Li, Pierre Reinprecht, Bruce Wittmann, Jennifer L Kennemur, Hayden Carder, Jason Yang, Theophile Lambert, Ryen O’Meara, Lukas Radtke, Ziyang Qin, Sabine Brinkmann-Chen, Frances Arnold, and Ariane Mora. Enzyme engineering database (enzengdb): a platform for sharing and interpreting sequence–function relationships across protein engineering campaigns. Nucleic Acids Research, page gkaf1142, December 2025. ISSN 1362-4962. doi: 10.1093/nar/gkaf1142.

[40] Gemini Gemini Team, Rohan Anil, Sebastian Borgeaud, Jean-Baptiste Alayrac, Jiahui Yu, Radu Soricut, Johan Schalkwyk, Andrew M Dai, Anja Hauth, Katie Millican, et al. Gemini: a family of highly capable multimodal models. arXiv *preprint arXiv:2312.11805*, 2023.

[41] Esther Heid, Daniel Probst, William H Green, and Georg KH Madsen. Enzymemap: curation, validation and data-driven prediction of enzymatic reactions. Chemical Science, 14(48):14229–14242, 2023.

[42] William JF Rieger, Mikael Boden, Frances Arnold, and Ariane Mora. Squidly: Enzyme Catalytic Residue Prediction Harnessing a Biology-Informed Contrastive Learning Framework. January 2026. doi: 10.7554/elife.108186.2. URL 10.7554/eLife.108186.2.

[43] Vignesh Ram Somnath, Matteo Pariset, Ya-Ping Hsieh, Maria Rodriguez Martinez, Andreas Krause, and Charlotte Bunne. Aligned diffusion schrödinger bridges. In Uncertainty in Artificial Intelligence, pages 1985–1995. PMLR, 2023.

[44] Greg Landrum, Paolo Tosco, Brian Kelley, Ricardo Rodriguez, David Cosgrove, Riccardo Vianello, Peter Gedeck, Gareth Jones, Eisuke Kawashima, Dan Nealschneider, et al. rdkit/rdkit: 2025_03_1 (q1 2025) release. Zenodo, 2025.

[45] Ladislav Rampášek, Mikhail Galkin, Vijay Prakash Dwivedi, Anh Tuan Luu, Guy Wolf, and Dominique Beaini. Recipe for a general, powerful, scalable graph transformer, 2023. URL https://arxiv.org/abs/2205.12454.

[46] UniProt Consortium. Uniprot: a worldwide hub of protein knowledge. Nucleic acids research, 47(D1):D506–D515, 2019.

[47] Christiam Camacho, George Coulouris, Vahram Avagyan, Ning Ma, Jason Papadopoulos, Kevin Bealer, and Thomas L Madden. Blast+: architecture and applications. BMC bioinformatics, 10(1):421, 2009.

[48] Kazutaka Katoh, Kazuharu Misawa, Kei-ichi Kuma, and Takashi Miyata. Mafft: a novel method for rapid multiple sequence alignment based on fast fourier transform. Nucleic acids research, 30(14):3059–3066, 2002.

[49] Xiaohong Ji, Zhen Wang, Zhifeng Gao, Hang Zheng, Linfeng Zhang, Guolin Ke, and Weinan E. Uni-mol2: Exploring molecular pretraining model at scale, 2024. URL https://arxiv.org/abs/2406.14969v2.

